# Augmentation of Antibacterial Activity in Mesenchymal Stromal Cells Through Systems-Level Analysis and CRISPR-mediated Activation of CD14

**DOI:** 10.1101/2020.10.14.338020

**Authors:** Matthew P. Hirakawa, Nikki Tjahjono, Yooli K. Light, Prem Chintalapudi, Kimberly S. Butler, Steven S. Branda, Raga Krishnakumar

**Affiliations:** Systems Biology Department, Sandia National Laboratories, Livermore, CA. 94551, USA; Molecular & Microbiology Department, Sandia National Laboratories, Albuquerque, NM. 87185, USA; Molecular & Microbiology Department, Sandia National Laboratories, Livermore, CA. 94551, USA

## Abstract

Mesenchymal stromal cells (MSCs) have broad-ranging therapeutic capabilities, however MSC use is confounded by cell-to-cell heterogeneity, and source-to-source phenotypic inconsistencies. We utilized a systems-based approach to compare MSCs which displayed different capacity for antibacterial activity. Although MSCs from both sources satisfied traditional MSC-defining criteria, comparative transcriptomics and quantitative membrane proteomics demonstrated two unique molecular profiles. The antibacterial MSCs respond rapidly to bacterial lipopolysaccharide (LPS) and have elevated levels of the LPS co-receptor CD14. CRISPR-mediated overexpression of endogenous CD14 in non-antibacterial MSCs resulted in faster LPS response and enhanced antimicrobial activity. Single-cell transcriptome profiling of CD14-activated MSCs revealed uniform enhancement of LPS response kinetics, and a shift in the ground state of these MSCs. Our results demonstrate that systems-level analysis can reveal critical molecular targets to optimize desirable properties in MSCs, and that overexpression of CD14 in these cells can shift their state to be more responsive to future bacterial challenge.

## INTRODUCTION

Mesenchymal stromal cells (MSCs) are a group of multipotent and phenotypically plastic cells that have enormous potential for use in diverse clinical applications ^1–3^. A major roadblock inhibiting the progression of MSCs for clinical use is their extensive cell-to-cell heterogeneity and phenotypic variation that constrains experimental reproducibility and behavior prediction ^1,3^. MSC phenotype is strongly dependent on their origin, as cells isolated from different donors exhibit distinct immunogenic and therapeutic properties based on the health and age of the donor, making the efficacy of these cells unpredictable ^4–7^. Additionally, MSCs can be isolated from virtually every post-natal body site, but MSCs from these different body sites exhibit site-specific phenotypes with distinct proliferative lifespans, immunomodulatory properties, epigenetic markers, transcriptional profiles, and cell surface proteins ^8–16^. Even populations of MSCs isolated from a single site of a donor are heterogenous, and clonal propagation of a single cell leads to intra-colony heterogeneity ^17^. Although MSC heterogeneity has made it difficult to define and reproducibly characterize these cells, it is likely this same cellular variation that enables MSCs to exhibit such extensive phenotypic diversity. A better understanding of the molecular mechanisms regulating specific phenotypes, and methods to homogenize and regulate MSC behavior, are crucial steps towards the utilization of MSCs in future therapeutic applications.

A significant threat to global human health that requires the development of new and alternative therapies is the emergence of drug resistant microorganisms. MSCs are intriguing candidates as countermeasures against microbial infections because they are capable of directly inhibiting microbial growth and modulating host immune responses by regulating immune cell localization at sites of infection through secretion of pro- and anti-inflammatory cytokines ^18,19^. With regard to direct antimicrobial activity, MSCs have been shown to inhibit the growth of bacterial and fungal pathogens both *in vitro* and *in vivo* ^19–23^. These antibacterial effects of MSCs are, in part, due to the secretion of antimicrobial peptides including the cathelicidin LL-37, β-defensin 2, lipocalin-2, and hepcidin, and the administration of MSCs has been shown to promote resolution of microbial infection in animal models ^20–25^. MSCs have the potential for broad-acting cell-based antimicrobial therapy, however the molecular basis for antimicrobial activity, and conditions to maintain MSC populations of uniform phenotype are not fully understood. Therefore, strategies for generating MSC populations of predictable behavior will be required for their safe and reliable use in clinical applications.

One such strategy to direct, or enhance, MSC properties is through “priming” these cells by specifically altering their environment to direct their phenotypic output towards desired therapeutic functions ^1,26,27^. For example, exposure of MSCs to hypoxia and serum deprivation (a condition that mimicks *in vivo* administration of these cells) leads to metabolic and lipidomic reprogramming, as well as increased production of exosomes packed with immunomodulatory metabolites ^28–30^. In another example, priming MSCs with cytokines can direct large-scale changes to the MSC proteome and secretome that impact inflammation and angiogenesis ^31,32^. Importantly, these priming stimuli *in vitro* can create enduring alterations in MSC phenotypes which persist even upon transition to a new environment, including *in vivo* contexts; this suggests that “priming” may be an effective strategy for manipulating MSC behavior for therapeutic purposes ^26,33^. Priming MSCs using Toll-like receptor (TLR) agonists are particularly notable, as stimulation of these receptors leads to diverse phenotypic alterations to MSCs ^34–38^. There are at least 13 TLRs encoded by mammalian genomes, each specialized to recognize specific pathogen associated molecular patterns (PAMPs), together enabling the host to respond to presence of viral, bacterial and fungal pathogens ^39^. The expression of TLRs in MSCs are origin- and context-dependent, however stimulation of these receptors can modulate diverse biological processes in these cells ^34–38^. For example, stimulation of TLR3 causes MSCs to exhibit enhanced migration, upregulates immunosuppressive traits, and provides therapeutic benefits for colitis ^40,41^. Additionally, systemic administration of TLR3-primed MSCs in animals with chronic bacterial infections can promote dramatic reduction in bacterial burden at sites of infection and promote wound healing ^42^. Interestingly, while TLR3 priming tends to drive MSCs towards an immunosuppressive state, TLR4 priming shifts MSCs into a more proinflammatory state ^40^. TLR4 priming, by exposing MSCs to bacterial lipopolysaccharide (LPS), causes widespread transcriptional changes primarily regulated by NF-κB and IRF1, and leads to the upregulation of genes involved with inflammatory response and chemotaxis ^43^. Interestingly, LPS-primed MSCs have also been demonstrated to have enhanced antibacterial properties due to the upregulation of antimicrobial peptides ^20,21,44^. Additionally, TLR4 has been identified as a critical receptor that promotes antibacterial activity in MSCs, and knockdown of this protein reduces the ability of MSCs to inhibit *E. coli* growth ^21^. Although TLR4 priming leads to MSCs with enhanced antibacterial activity, the underlying genetic mechanisms regulating this antibacterial state, and tools to identify and homogenize populations of antimicrobial MSCs, have yet to be examined.

MSCs are most frequently identified and defined based on the presence of MSC-specific cell-surface markers; however, cells expressing equivalent markers frequently exhibit distinct phenotypes and biological properties (e.g. MSCs from different body sites) ^8,16,45,46^. Extensive efforts have been made to link MSC subpopulations from specific sources or with distinct phenotypes to cell surface marker profiles, often using fluorescence-activated cell sorting (FACS) to purify MSCs with specific traits ^47–49^. However, as mentioned above, even cells derived from a single source can contain extensive heterogeneity. Recent advances in single-cell sequencing technologies enable higher resolution characterization of cellular subtypes and population diversity. These techniques are particularly informative to investigate molecular signatures of MSCs exhibiting a particular trait, or molecular changes after MSCs are exposed to priming conditions. Although at the single-cell level there exists significant cell-to-cell transcriptional heterogeneity, priming strategies have been shown to promote population-homogenizing effects that unify cellular behavior in inherently heterogenous MSCs ^50–53^. Therapeutic application of MSCs depends on the predictability of their *in vivo* phenotype, which requires, in turn, effective control strategies such as priming. New genetic engineering tools, such as CRISPR-based gene activation or repression, may enable novel approaches for optimizing control strategies. Such optimization also requires an adequate systems biology understanding of the developmental, transcriptomic and proteomic states of heterogenous MSC populations.

In this work, MSCs from the bone marrow of two different mouse strains (C57BL/6 and BALB/c) were examined for their ability to inhibit bacterial growth, with C57BL/6 MSCs (C57-MSCs) showing significantly stronger antimicrobial activity compared with BALB/c MSCs (BALB-MSCs). Using a combination of molecular profiling strategies, we identified striking differences between these two MSC subtypes regarding their ability to sense and respond to bacteria. Based on these analyses, we focused on the LPS co-receptor CD14, which was expressed at higher levels in C57-MSCs. Using CRISPR-activation (CRISPRa) to turn on expression of CD14 in BALB-MSCs, we were able to enhance their ability to inhibit bacterial growth, even in the absence of a priming agent. We further characterized these CD14 activated MSCs at the single-cell transcriptional level and demonstrated that CD14 expression in BALB-MSCs potentiates a more rapid response to bacteria-derived LPS. Overall, we demonstrate that a systems biology approach can reveal critical targets for enhancing specific MSC phenotypes, and that CRISPR-based gene modulation is an effective strategy to engineer MSCs with potential therapeutic properties.

## RESULTS

### MSCs exhibit source-dependent antibacterial phenotypes

The cells used in this study were commercially available bone marrow derived MSCs isolated from C57BL/6 or BALB/c mice and were examined for their expression of established mouse MSC surface markers and their ability to differentiate ^54^. We first examined the expression levels of mouse MSC markers in C57-MSCs and BALB-MSCs and observed that both cell types exhibited similar levels of these proteins using immunostaining followed by flow cytometry (**Supplementary Figure 1**). These cells were also inspected for their ability to differentiate into adipocytes and osteoblasts, and both C57-MSCs and BALB-MSCs could differentiate under adipogenic and osteogenic inducing conditions **(Supplementary Figure 2)**^55^. Using these traditional metrics to define cell identity, both C57-MSCs and BALB-MSCs could be considered the same cell type.

Although these C57-MSCs and BALB-MSCs are nominally the same cell subtype, we observed them to exhibit strikingly different phenotypes when co-cultured with bacteria. Here, we tested the ability of MSCs to limit bacterial growth by co-culturing the MSCs with *E. coli* K-12 MG1655 (referred to as *E. coli* henceforth) *in vitro* (**Figure 1A, B**). *E. coli* were also independently co-cultured with 3T3 cells, a mouse embryonic fibroblast (MEF)-derived cell line, to serve as a control. After 6 hours of growth with 3T3 cells, the *E. coli* had grown from an initial concentration of 1 × 10^3^ CFU/ml to a final concentration of 1.1 × 10^6^ CFU/ml. When *E. coli* were cultured with C57-MSCs, the final bacterial concentration was reduced ~3-fold as compared to 3T3s (p=0.024), indicating that the C57-MSCs were capable of inhibiting bacterial growth. Surprisingly, the opposite result was observed when *E. coli* were co-cultured with BALB-MSCs, and resulted in a ~3-fold increase in bacterial CFUs when compared to 3T3s (p=0.0024), and a ~10-fold increase compared to C57-MSCs (p=8.45×10^−5^). Together, these data demonstrate that while MSCs isolated from different sources share hallmark features of MSCs, their antibacterial properties are distinct, which may have important implications regarding the efficacy of MSCs as potential therapy to treat bacterial infections.

**Figure 1.**
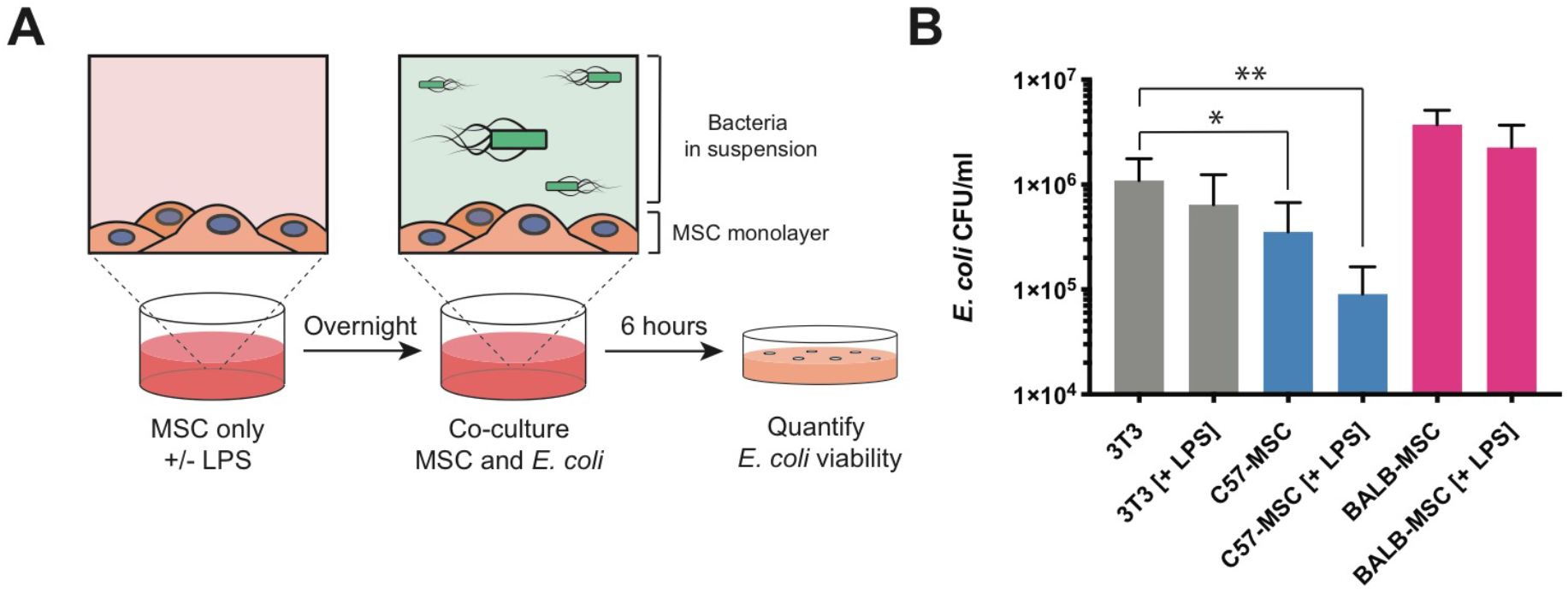
MSCs from different sources have distinct antibacterial properties. **(A)** Cartoon depicting the antibacterial assays used in this study. MSCs were incubated overnight with or without lipopolysaccharide (LPS) from *E. coli* (100 ng/ml) and were then co-cultured with *E. coli* strain K12 for 6 hours. After 6 hours of co-culture, *E. coli* abundance in the media was quantified using colony forming unit (CFU) assays. **(B)** Quantification of *E. coli* CFUs following 6-hour co-culture with different cell types +/− LPS treatment (n = at least 5 biological replicates per condition; * *P* <0.05; ** *P* < 0.005; error bars = standard deviation).

Priming MSCs via exposure to bacteria, or bacteria-derived LPS, has been shown to further enhance their antibacterial properties ^20,21,44^. Here, C57-MSCs and BALB-MSCs were primed with the TLR agonist LPS isolated from *E. coli* 055:B5 and tested for their ability to inhibit bacterial growth (**Figure 1A, B**). Pretreatment of control 3T3 cells with LPS did not result in a significant change when compared to untreated 3T3 cells (p=0.25). In contrast, C57-MSCs pre-treated with LPS further reduced *E. coli* CFUs ~12-fold when compared to the *E. coli* abundance in the presence of 3T3s (p=0.024), and ~4-fold reduction when compared to untreated C57-MSCs (although this reduction was not quite statistically significant, p=0.06). When BALB-MSCs were primed with LPS, there was a slight reduction in *E. coli* CFUs (~1.6-fold less) when compared to untreated BALB-MSCs, however this change was not statistically significant (p=0.14). These results prompted us to investigate the molecular mechanisms contributing to the difference in C57-MSC vs BALB-MSC antibacterial activity.

### C57-MSCs exhibit more rapid transcriptional response to LPS stimulation than BALB-MSCs

LPS exposure triggers a number of signaling pathways in mammalian cells leading to the nuclear translocation of cytoplasmic sequestered transcription factors, including the NF-κB family of transcription factors, that are key regulators of antimicrobial activity and inflammation ^56–59^. We hypothesized that the absence of antibacterial activity exhibited by BALB-MSCs may be the result of an attenuated or delayed recognition of bacterial exposure when compared to the C57-MSC response. To examine temporal responses to LPS stimulation, C57-MSCs and BALB-MSCs were treated with LPS and nuclear translocation of the NF-κB subunit p65 was assayed using immunostaining followed by quantitative high-throughput microscopy (**Figure 2A**). In these assays, C57-MSCs responded to LPS stimulation with a peak nuclear p65 localization (~85% of cells analyzed) occurring at 0.5 hrs post-exposure. In contrast with the C57-MSCs, only ~27% of BALB-MSCs displayed nuclear p65 at the 0.5 hr timepoint (p=3×10^−6^). Instead, the BALB-MSCs exhibited peak nuclear p65 staining (~90% of cells) occurring at 2 hours post-exposure, indicating that BALB-MSCs were somehow slower to transduce the LPS signal from the cell surface into the nucleus when compared to C57-MSCs.

**Figure 2.**
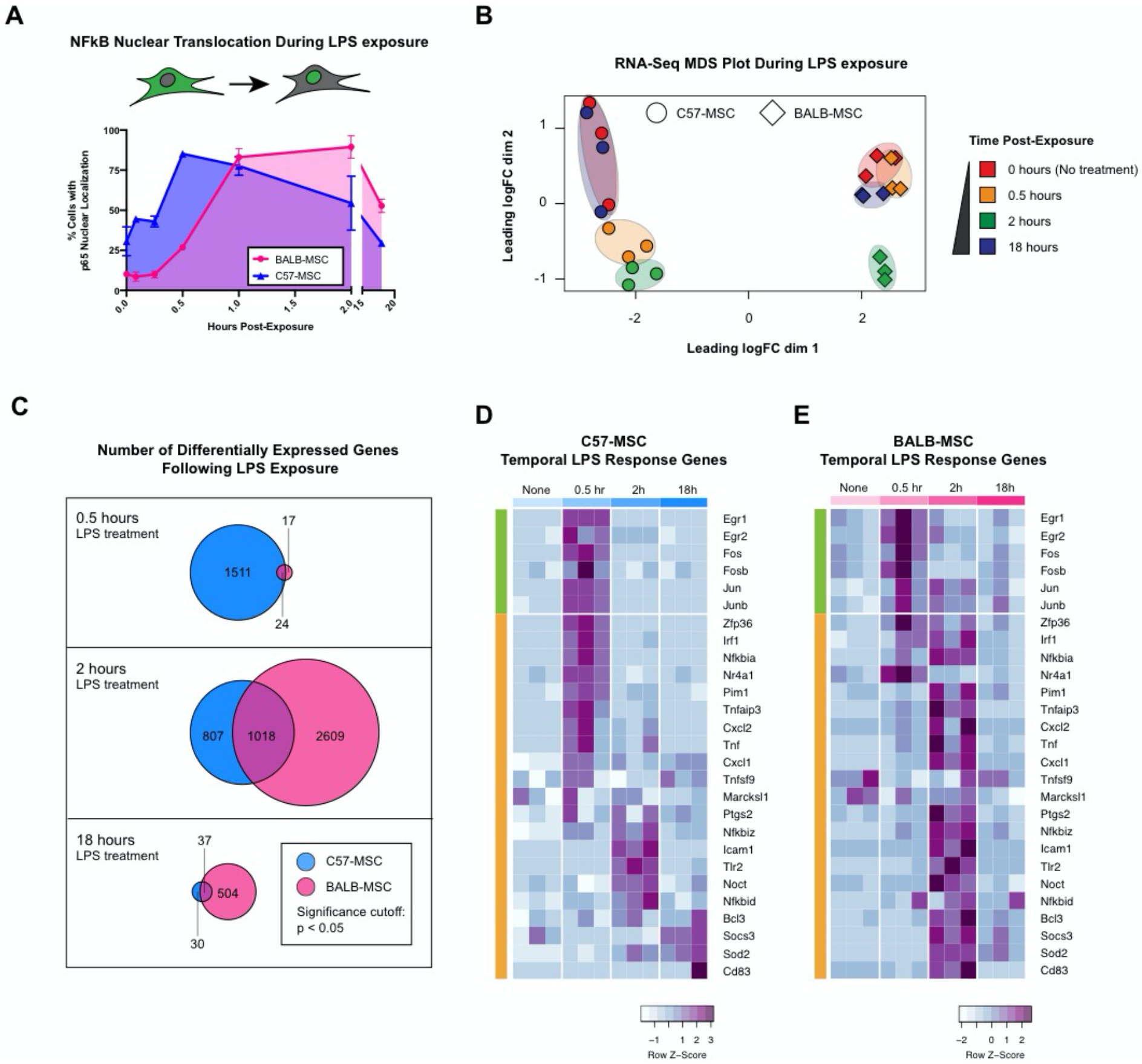
Differential response rates to LPS between MSC sources. **(A)** Quantification of NF-κB nuclear translocation assays using p65 staining and high-throughput imaging (>1000 cells analyzed per replicate; n=3 biological replicates; error bars represent standard deviation). **(B)** MDS plot depicting transcriptional profiles of C57-MSCs and BALB-MSCs at timepoints post LPS-exposure (0 hr, 0.5 hr, 2 hr, and 18 hr). **(C)** Numbers of differently expressed genes at different timepoints post LPS-exposure compared to corresponding untreated cells. **(D, E)** Heatmap depicting temporal expression of immediate-early genes (green bars) and early LPS response (orange bars) in C57-MSC and BALB-MSC at timepoints post LPS-exposure (immediate-early genes and early LPS response genes were previously defined ^60–62^).

Because we observed C57-MSCs and BALB-MSCs to exhibit distinct NF-κB nuclear translocation kinetics in response to LPS, we hypothesized that rates of transcriptional change would also differ in these two cell types. To examine patterns of gene expression following LPS exposure, population-level transcriptional profiling (RNA-seq) was performed on MSCs prior to LPS exposure, and at three timepoints following LPS exposure (0.5 hrs, 2 hrs, and 18 hrs). To visualize the relationships between MSC transcriptional profiles after LPS exposure, read-count matrices were plotted using multi-dimensional scaling (**Figure 2B**). This approach clearly separated the C57-MSCs from the BALB-MSCs along dimension 1 suggesting these two cell types have distinct transcriptional profiles. Changes in transcriptional profiles that corresponded to the time following LPS exposure were apparent along dimension 2. C57-MSCs collected at 0.5 hrs or 2 hrs post-exposure formed two distinct clusters, indicating that their transcriptomes at these timepoints differed from that of resting-state C57-MSCs. In contrast, C57-MSCs at 18 hrs post-exposure clustered very closely with their untreated counterparts, indicating that their transcriptome had returned to the resting state. Consistent with their slower response to LPS exposure as evident from NF-κB translocation kinetics, the BALB-MSC transcriptome at 0.5 hrs post-exposure closely resembled that of untreated BALB-MSCs. By 2 hrs post-exposure, the BALB-MSC transcriptome was clearly distinct from that of untreated BALB-MSCs, clustering in a similar position as C57-MSCs along dimension 2; and by 18 hrs post-exposure returned to the resting state (i.e., resembled the untreated BALB-MSC transcriptome). These results indicate that after exposure to LPS, the C57-MSC transcriptome is rapidly altered, whereas the BALB-MSC transcriptome shows delayed response.

We next performed differential gene expression analysis to identify genes that had significant changes in expression after LPS exposure (**Figure 2C**). We found that 1535 genes were differentially expressed in C57-MSCs by 0.5 hrs post-exposure, with the majority (926 genes) being upregulated. Enriched functional gene ontology (GO) categories of these upregulated genes included regulation of adaptive immune response, and cortical cytoskeleton organization. Of particular note, the genes displaying greatest fold change included those encoding chemokines Cxcl1 and Cxcl2, the NF-κB inhibitor Nfkbia, and an RNA-binding protein involved with early LPS response (Zfp36). These results indicate that by 0.5 hrs post-exposure, the C57-MCSs had mounted a multifaceted response including gene programs relevant for combating infection. In contrast, only 41 genes were differentially expressed in BALB-MSCs by 0.5 hrs post-exposure. The small number of differentially expressed genes was not sufficient to support GO category enrichment analysis; however, several of the genes are implicated in early LPS response and were similarly differentially expressed in C57-MCs by 0.5 hrs post-exposure (Jun, Cxcl1, Nfkbia, Tnfaip, Zfp36, and Ier3).

We further examined a subset of previously characterized early LPS response genes to better understand the timing of activation of these genes in our MSCs. These early response genes include immediate-early genes (IEGs) that are conserved across cell types and are rapidly activated by diverse environmental stimuli ^60,61^. Following IEG activation, primary LPS response genes are activated that are more specific to this stimulus ^60,62^. Consistent with the rapid activation of IEGs following LPS stimulation, both C57-MSCs and BALB-MSCs displayed robust up-regulation of the conserved IEGs Egr1, Egr2, Fos, Fosb, Jun, and Junb by 0.5 hrs post-exposure (**Figure 2D, E**). However, the expression of primary LPS response genes varied with MSC source, with the majority showing peak expression at 0.5 hrs post-exposure in C57-MSCs vs 2 hrs post-exposure in BALB-MSCs. Together, these results indicate that by 0.5 hrs post-exposure, the C57-MSCs had already progressed into an LPS specific response, whereas the BALB-MSCs were only beginning to undergo changes in gene expression involved with generic stimulus exposure.

Finally, we examined upregulated genes at 18 hrs post-exposure, as this is the timepoint at which MSCs were challenged with *E. coli* in our antibacterial assays. At this timepoint, both C57-MSCs and BALB-MSCs were more similar to untreated cells as compared to their transcriptional profiles at 2 hrs post-exposure. However, there were notable genes with elevated expression that have previously been implicated in innate immune response or antibacterial activity. In particular, C57-MSCs showed increased expression of the genes encoding chemokines Cxcl1 and Cxcl5, nitric oxide synthase Nos2, a serine protease inhibitor Serpina3, and the proinflammatory cytokine Il-6. BALB-MSCs exhibited more differentially expressed genes (541), with those upregulated (376) tending to belong to GO categories for LPS-mediated signaling and regulation of inflammatory response. Similar to C57-MSCs, among the most upregulated genes at 18 hrs post-exposure were those involved with innate immune responses or direct antibacterial activity; these included Nos2, Il-6, Cxcl1, Cxcl5, the iron-sequestering protein lipocalin-2 (Lcn2) and complement component 3 (C3). Thus, by 18-hrs post-LPS exposure, the overall transcriptomes of C57-MSCs and BALB-MSCs very closely resembled their unstimulated counterparts, however both cell types exhibited elevated expression of a handful of genes that are involved with recruitment of immune cells and antibacterial activity.

### C57-MSC vs BALB-MSC expression of genes mediating LPS recognition

Because C57-MSCs responded more quickly to LPS and were capable of mounting a more robust antibacterial response than BALB-MSCs, we hypothesized that the baseline transcriptional state of C57-MSCs may be different from that of BALB-MSCs to allow them to respond more quickly to bacterial exposure. To test this hypothesis, we used RNA-seq to examine the transcriptomes of unstimulated C57-MSCs and BALB-MSCs. We additionally performed RNA-seq on two fibroblast cell types for comparison: primary mouse embryonic fibroblasts (MEFs) and primary mouse dermal fibroblasts (MDFs) from C57BL/6 mice. To define the relationships between cell types, transcriptional profiles were first visualized using principal component analysis (PCA) (**Figure 3A**). Using this approach, the C57-MSCs and BALB-MSCs clustered separately along PC1, but both diverged similarly from the fibroblasts along PC2. The C57-MSCs and BALB-MSCs shared 523 upregulated genes when compared to both fibroblasts (**Supplementary Figure 3A**). These genes were associated with multiple GO-term categories ascribed to innate immune response and could be summarized by a single statistically significant GO-slim category “defense response” (**Supplementary Figure 3B**). Although we do not further characterize them in this work, these genes found to be specifically up-regulated in MSCs could serve as additional markers for identification or isolation of MSCs.

**Figure 3.**
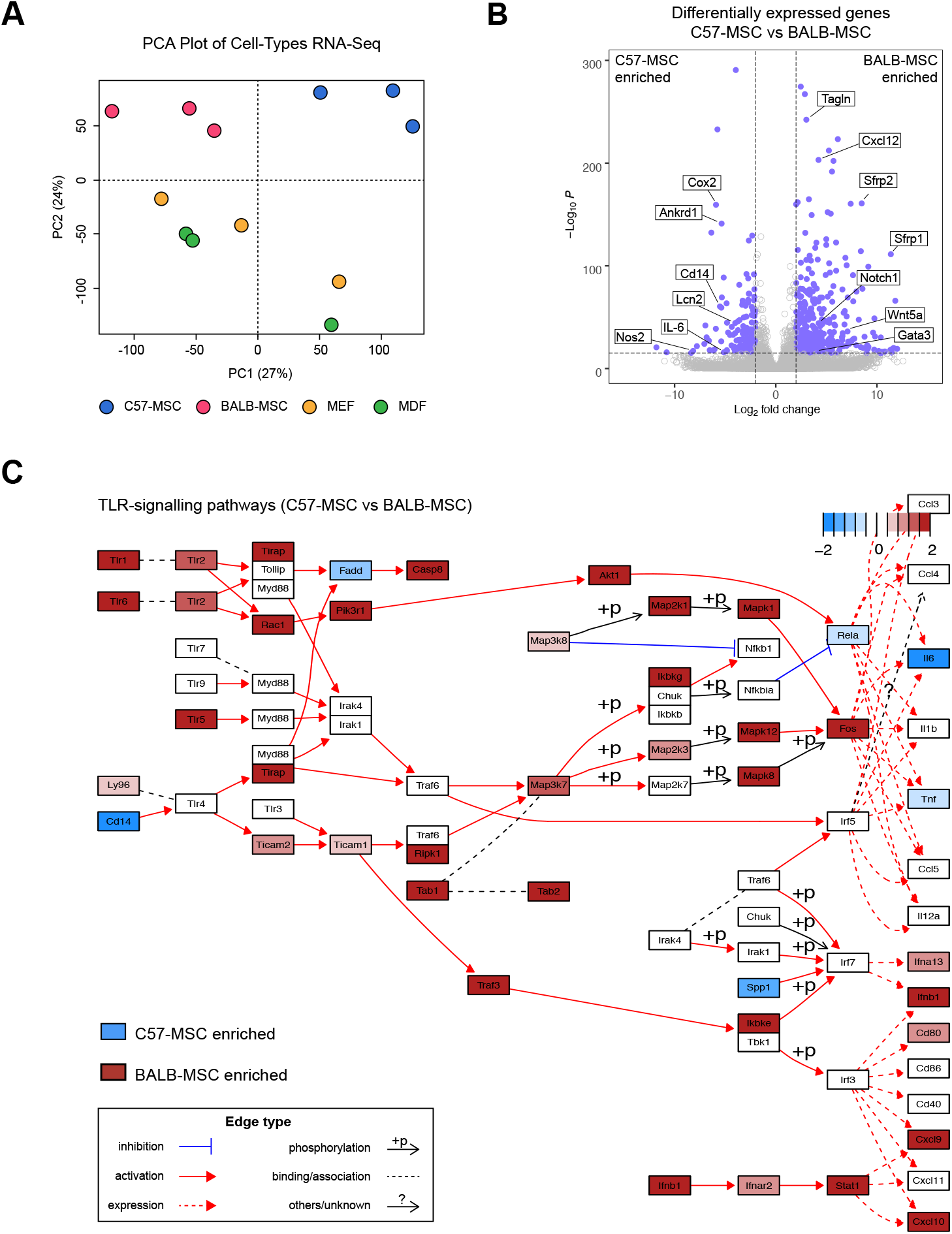
Comparative transcriptional analyses of MSC types using RNA-seq. **(A)** PCA plot depicting transcriptional profiles using RNA-seq of C57-MSCs, BALB-MSCs, MEFs and MDFs under standard growth conditions. **(B)** Volcano plot depicting differentially expressed genes between C57-MSCs and BALB-MSCs and highlighting notable genes of interest (blue dots represent genes with fold change > 2 and adjusted P-value < 10^−16^). **(C)** Pathway analysis visualization of comparative gene expression between C57-MSCs (blue) and BALB-MSCs (red) of TLR-signaling pathways.

To further highlight the differences between C57-MSCs and BALB-MSCs at the transcriptional level, RNA-seq data from the two MSC types were directly compared through differential expression analysis. We identified 662 genes that were upregulated in C57-MSCs relative to BALB-MSCs; and 1324 genes upregulated in BALB-MSCs relative to C57-MSCs. These genes that are specifically enriched in C57- or BALB-MSCs were then used in GO-term analysis to identify over-represented biological functions (**Supplementary Figure 4)**. Among the enriched GO-categories in C57-MSCs was the response to LPS, which supported our functional observations that these cells respond more quickly to LPS exposure.

We next identified the genes exhibiting the highest levels of differential expression between baseline C57-MSCs and BALB-MSCs to determine whether they could provide additional insight into differences in antibacterial activity. Interestingly, among the most upregulated genes in C57-MSCs included those involved with inflammation and antibacterial responses, including Ptgs2 (Cox2), Lcn2, Nos2, Il-6, and Cd14 (**Figure 3B**). These genes are all known to be regulated by NF-κB or direct activators of the NF-κB pathway, which in turn is closely tied to TLR-mediated pathogen recognition ^63–70^. Higher levels of secreted Il-6 protein were also observed in C57-MSCs before and after LPS-treatment through cytokine profiling experiments providing some support linking RNA-seq data with protein expression levels (**Supplementary Figure 5**). Among the most upregulated genes in BALB-MSCs were those involved with development and cell polarization (e.g., Gata3, Notch1, Wnt5a, Sfrp1 and Sfrp2) which corresponded with many of the GO-term categories enriched in these cells. Because C57-MSCs was more responsive to LPS, we hypothesized that these C57-MSCs would display higher expression levels of genes participating in TLR signaling pathways as compared to BALB-MSCs. However, using the pathway analysis tool Pathview ^71^ we unexpectedly observed that the majority of TLR pathway components were more highly expressed in BALB-MSCs (**Figure 3C**). However, a striking exception in these pathways was the elevated expression level of the LPS co-receptor CD14 in C57-MSCs. Because we had observed C57-MSCs to respond more rapidly to LPS stimulation and expressed higher levels of CD14, we examined this gene further in subsequent sections of this work.

### Comparative Membrane Proteomics Between Cell Types

To further investigate molecular differences between C57-MSCs and BALB-MSCs, we utilized a proteomics-based approach to characterize protein levels between these cell types using liquid chromatography-mass spectrometry (LC-MS). We focused on membrane proteins, as these would likely mediate direct interaction with bacteria and initiate downstream signaling. To visualize membrane protein profiles, label-free quantification (LFQ) intensities were plotted using PCA and hierarchical clustering to identify sample relatedness (**Fig. 4A** **and** **4B**). MEFs and MDFs clustered closely together, indicating that these cell types have similar membrane protein compositions compared to the MSCs. The C57-MSCs and BALB-MSCs clustered together along PC1, away from the MEFs and MDFs; but diverged from each other along PC2. Using differential protein expression analysis, we identified 1,003 proteins that showed significant differences in abundance between the MSC types, with 438 more abundant in C57-MSCs, and 565 more abundant in BALB-MSCs. We next utilized GO analysis to determine if these differentially expressed proteins could point to any specific biological functions that may provide insight into their differences in antibacterial activity (**Supplementary Figure 6**). Among the most enriched GO categories in C57-MSCs included proteins involved with insulin binding and cell adhesion binding, while BALB-MSCs were enriched for TAP binding and aminopeptidase activity. We also examined the most differentially expressed membrane proteins between the two MSC types to inspect for proteins that may be associated with antibacterial phenotypic differences (**Figure 4C**). Notably, among the most enriched proteins in C57-MSCs was the LPS receptor CD14, consistent with RNA-seq data presented in Figure 3. CD14 protein expression was also observed in C57-MSCs, but not BALB-MSCs, using immunostaining followed by flow cytometry (**Supplementary Figure 7**). Among the most enriched proteins in BALB-MSCs were a membrane metallo-endopeptidase (Mme, also known as neprilysin) and ephrin type-B receptor, Ephb2. Comparing CD14 levels across all four cell types analyzed revealed that only C57-MSCs expressed high levels of CD14 protein (**Figure 4D**), suggesting that this protein may be critical for mounting an efficient LPS response and promoting antibacterial properties. Together, through our functional LPS response assays, comparative transcriptomic and proteomic analyses, we hypothesized that BALB-MSCs may have attenuated antibacterial properties due to a lack of CD14. In the remaining sections, we tested this hypothesis by upregulating endogenous CD14 using CRISPR tools and examined the LPS-response and antibacterial properties of these engineered BALB-MSCs.

**Figure 4.**
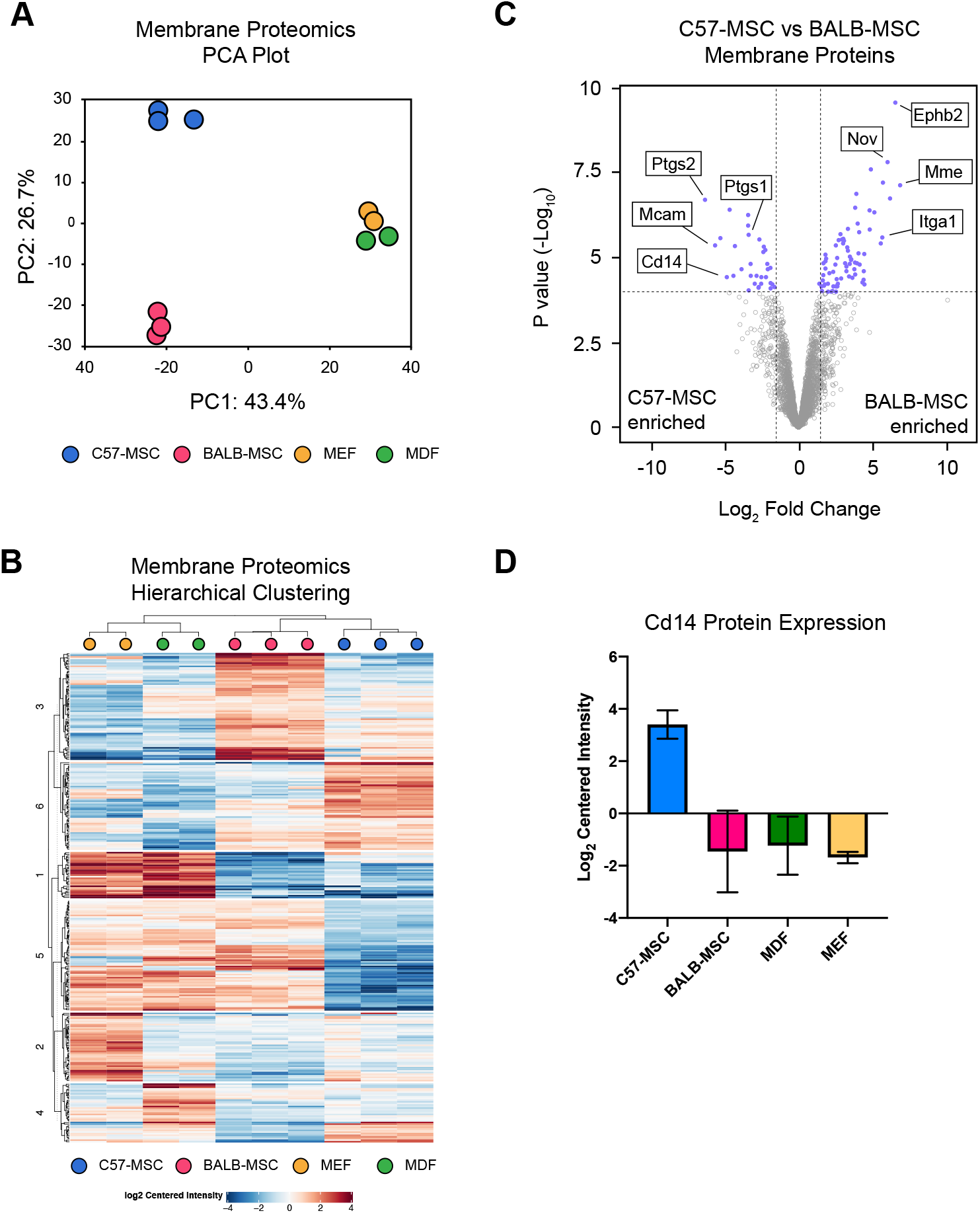
Characterization of cell membrane protein differences between cell types using quantitative proteomics. MSCs and fibroblasts were grown in standard culture conditions and membrane proteins were analyzed using LC-MS/MS. **(A)** PCA plot depicting membrane protein profiles of MSC and fibroblast cell types (n=3 biological replicates for MSCs, and n=2 biological replicates for MEF and MDF). **(B)** Heatmap utilizing hierarchical clustering showing expression level differences of membrane proteins. **(C)** Volcano plot highlighting significantly differentially expressed proteins between C57-MSCs and BALB-MSCs and notable proteins of interest (blue dots represent proteins with fold change > 2 and adjusted P-value < 10^−4^). **(D)** Protein expression levels of the LPS-receptor CD14 across the cell types examined in this study (n = 3 biological replicates, error bars = standard deviation).

### Overexpression of endogenous CD14 enhances antibacterial properties in non-antibacterial MSCs

CD14 facilitates LPS recognition by TLR4, significantly enhancing the response to LPS in mammalian cells and in mice ^72,73^. In this study, we observed BALB-MSCs and C57-MSCs to exhibit distinct LPS response kinetics, and through transcriptional and proteomic profiling we found CD14 was differentially regulated between these two cell types. We therefore hypothesized that the upregulation of CD14 could augment the attenuated antibacterial activity in BALB-MSCs. To test this hypothesis, BALB-MSCs were engineered to overexpress endogenous *Cd14*, through the use of CRISPR-mediated activation (CRISPRa). Although differential expression of *Tlr4* was not observed between cell types, overexpression of this gene was also examined independently here as it functions in conjunction with *Cd14* and has a well-established role in propagation of LPS signaling ^39,57^. First, BALB-MSC-CRISPRa were constructed using lentiviral vectors to stably express the CRISPRa Synergistic Activation Mediator (SAM) system ^74^. Second, BALB-MSC-CRISPRa were tested for overexpression of target genes using six sgRNAs that localized to different distances from the *Cd14* or *Tlr4* transcription start site (TSS). MSCs expressing different sgRNAs were tested for protein expression levels by immunostaining followed by flow cytometry (**Supplementary Figure 8**). The sgRNAs that led to the most robust protein expression were used for subsequent experiments.

To test how overexpression of *Cd14* in BALB-MSCs impacted their antibacterial activity, these cells were co-cultured with *E. coli* and bacterial growth measured using CFU assays (**Figure 5A**). When *E. coli* was co-cultured with wildtype BALB-MSCs for 6 hours, the bacterial density reached 3.8 × 10^6^ CFU/ml. Similarly, when *E. coli* was co-cultured with BALB-MSCs expressing non-targeting control sgRNAs (BALB-MSC-CRISPRa-Scrambled), *E. coli* reached a final concentration of 3.3 × 10^6^ CFU/ml which was not significantly different from the wildtype control (p = 0.65). In contrast, when *E. coli* was grown with BALB-MSC-CRISPRa-CD14, we observed a ~45% reduction in bacterial growth compared to scrambled sgRNA control cells (p = 0.026). Overexpression of *Tlr4* in BALB-MSCs similarly trended towards reduced bacterial growth, although these results were not significant as there was a large deviation between replicates (p = 0.11) (**Supplementary Figure 9A**). These results demonstrate that the upregulation of endogenous *Cd14* can significantly improve antibacterial properties in MSCs, even in the absence of LPS priming.

**Figure 5.**
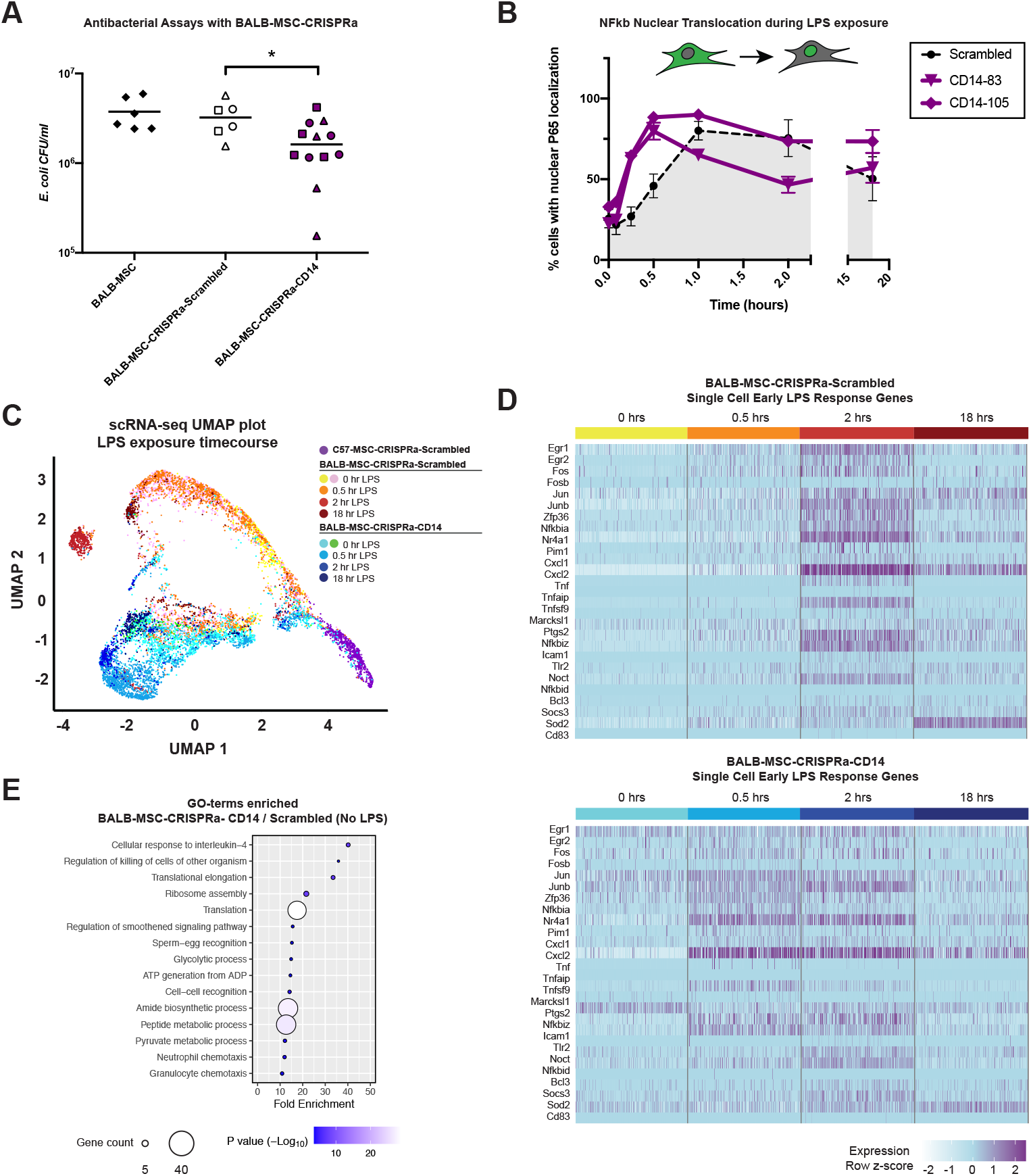
Antibacterial and single-cell transcriptional analyses of BALB-MSCs overexpressing endogenous CD14 via CRISPRa. **(A)** *E. coli* CFUs from antibacterial assays using wildtype and BALB-MSC-CRISPRa cells (*, P < 0.05). Three different sgRNAs were tested for CRISPRa cells and individual sgRNAs are represented with different shapes (For scrambled sgRNAs: circle = non-targeting_control_1, square = non-targeting_control_2, triangle = non-targeting_control_3; For CD14 sgRNAs: circle = CD14-83, square = CD14-105, triangle = CD14-131). **(B)** Nuclear translocation of NF-κB during LPS exposure using p65 immunostaining and quantitative microscopy in BALB-MSC-CRISPRa cells expressing a scrambled sgRNA or CD14 sgRNAs (scrambled = non-targeting_control_1; n = at least 3 biological replicates per sample). **(C)** UMAP plot depicting single-cell transcriptional profiles of MSCs during LPS-exposure (scrambled sgRNA = non-targeting_control_1; CD14 sgRNA = CD14-83). Single-cell RNA-seq of BALB-MSC overexpressing CD14 during LPS-exposure was prepared using the 10x Genomics Chromium platform followed by Illumina sequencing. **(D)** Heatmap of early LPS response gene expression in 100 individual BALB-MSC-CRISPRa cells expressing either scrambled control sgRNA (top) or CD14 sgRNA (bottom). **(E)** GO-terms enriched in differentially expressed genes from BALB-MSC-CRISPRa-CD14 compared to BALB-MSC-CRISPRa-CD14 prior to LPS-exposure.

As previously discussed, LPS stimulation triggers the translocation of NF-κB from the cytoplasm into the nucleus. To test how overexpression of *Cd14* affects the rate of NF-κB nuclear translocation, engineered BALB-MSCs were treated with LPS and nuclear translocation of the p65 NF-κB subunit was quantified at various timepoints using immunostaining and quantitative high-throughput microscopy (**Figure 5B**). In this experiment, the BALB-MSC-CRISPRa-Scrambled control exhibited peak levels of nuclear NF-κB at 1-hour post-LPS exposure. Importantly, the overexpression of *Cd14* in these BALB-MSCs caused NF-κB translocation to occur more quickly, with cells exhibiting peak levels of nuclear NF-κB at 30 minutes post-exposure (p < 0.01). Interestingly, *Tlr4* overexpression did not significantly affect NF-κB translocation kinetics, as these cells were not different from cells expressing a non-targeting control sgRNA (**Supplementary Figure 9B**). This suggests that one mechanism by which CD14-overexpressing BALB-MSCs increased their antibacterial activity was through a more rapid response to bacterial exposure.

### Overexpression of CD14 in BALB-MSCs increases the rate of LPS response as measured by single-cell RNA-seq

In order to more closely examine the molecular properties and kinetics of LPS response in CD14-overexpressing BALB-MSCs, we performed single-cell RNA-seq using the 10X Genomics Chromium single-cell sequencing platform. BALB-MSC-CRISPRa-Scrambled and BALB-MSC-CRISPRa-CD14 were analyzed at 0 hrs, 0.5 hrs, 2 hrs and 18 hrs post-LPS exposure. Additionally, we examined C57-MSC-CRISPRa expressing a scrambled sgRNA in the absence of LPS as a control cell type for comparison. Using this approach, we observed that these BALB-MSC-CRISPRa-CD14 displayed accelerated LPS response kinetics when compared to those cells expressing a scrambled control. These results were observed using dimensional reduction clustering approaches, UMAP and tSNE, to visualize global transcriptional profiles of individual cells (**Figure 5C, Supplementary Figure 10B**) ^75,76^. Populations of the BALB-MSC-CRISPRa-SCRAMBLED cells separated from the untreated cells 2 hrs post LPS-exposure. These BALB-MSC-CRISPRa-SCRAMBLED cells did not exhibit a shift in transcriptional profiles at 0.5 hr post LPS-exposure as these two samples were highly overlapping using UMAP and tSNE clustering approaches. Alternatively, a clear separation of BALB-MSC-CRISPRa-CD14 cells was observed by 0.5 hrs post-LPS exposure when compared to their corresponding untreated cells, supporting the observation that these cells can respond more rapidly to LPS. This temporal LPS response was also observed using a heatmap depicting the expression levels of early LPS response genes (similar to that in Figure 2) by examining 200 individual cells per timepoint (**Figure 5D**). From this data, we observed cells expressing the scrambled guide to have highest expression levels of the majority of these genes at 2 hrs post LPS-exposure. Alternatively, BALB-MSCs overexpressing CD14 had elevated expression of many early LPS response genes 0.5 hrs post-LPS exposure, consistent with the more rapid NF-κB translocation observed in these cells.

### Single cell transcriptomic profiling of CD14 overexpressing MSCs reveals a shift in ground-state and increased population homogeneity

To further examine the molecular states of the various cell types and conditions, we again examined population structure using tSNE and UMAP ^75,76^. While the two representations are not identical, these clustering approaches converged on a number of interesting points. First, we observed C57-MSCs and BALB-MSCs cells distinctly clustered away from each other supporting our previous observation that these cell types have distinct molecular profiles (**Supplementary Figure 10C**). Second, we observed C57-MSCs clustered more tightly together when compared to BALB-MSCs suggesting that the C57-MSCs are a more transcriptionally homogenous population. These cell clustering approaches were supported by pairwise n-dimensional Euclidean distances between cells of the same sample, and a shared nearest neighbor approach which also demonstrated the BALB-MSC population was more transcriptionally heterogenous than C57-MSCs (**Supplementary Figure 10D,E**). Third, the overexpression of CD14 in BALB-MSCs homogenized their transcriptional profiles as observed by more unified clustering and smaller pairwise n-dimensional Euclidian distance measurements (**Supplementary Figure 10F,G**). Interestingly, LPS exposure decreased Euclidean distances between cells of the same sample suggesting that LPS-priming helped to homogenize transcriptional profiles in these cells. And finally, in the absence of LPS, BALB-MSC-CRISPRa-CD14 clustered away from BALB-MSC-CRISPRa-Scrambled, demonstrating that upregulation of the CD14 receptor shifts the ground state of these MSCs. To define these differences in ground state, we examined the differentially expressed genes between BALB-MSC-CRISPRa-Scrambled and BALB-MSC-CRISPRa-CD14 cells and characterized the functions of these genes using GO-term analysis (**Figure 5E**). Among the top GO-term categories enriched in BALB-MSC-CRISPRa-CD14 was “killing cells of other organisms”, which resulted from these cells having high expression of Cxcl1 and Cxcl5. Together, data suggests the overexpression of CD14 in BALB-MSCs shifts their ground state that is more amenable to elicit a rapid response to bacterial exposure.

## DISCUSSION

In this work, we investigated the ability of MSCs from different sources to inhibit bacterial growth, examined molecular variations between MSC sources, and utilized a systems level approach to identify genetic targets to improve antibacterial properties. We demonstrate that although MSCs from different sources may share similar cell surface markers, other phenotypes of these cells are not necessarily equivalent. This work highlights the source-dependent phenotypic variation of MSCs and demonstrates a strategy to enhance desirable therapeutic properties in these cells using CRISPR tools.

Our initial observations were consistent with previous studies that demonstrated priming MSCs with bacteria or bacterial LPS could enhance antibacterial activity in these cells ^20,21,44^. However, we observed that the response to LPS stimulation in C57- and BALB-MSCs exhibited differential temporal dynamics, and later found that these different LPS response rates may contribute to the difference in antibacterial activity between C57-MSCs and BALB-MSCs. Previous studies have demonstrated that mouse strain backgrounds, including C57BL/6 and BALB/c, can show dramatic differences in immune responses ^77,78^. Through our RNA-seq and proteomics experiments, we identified that C57-MSCs and BALB-MSCs exhibited notable differences in the LPS signaling pathway. Of particular interest was the striking upregulation of the LPS receptor CD14 at the RNA and protein levels in C57-MSCs when compared to BALB-MSCs. CD14 was originally identified as a marker of monocytes, and has since been shown to dramatically increase cell sensitivity to LPS ^73,79,80^. CD14 is a co-receptor that binds LPS monomers and delivers them to the TLR4-MD-2 complex, initiating a signaling program that activates potent transcription factors, including NF-κB, AP-1 and IRFs, which regulate immune responses ^56,70,81^. We therefore tested whether upregulation of CD14 in BALB-MSCs could improve the rate of LPS response in these cells and enhance antibacterial activity, similar to that of C57-MSCs. Here, we utilized CRISPRa technology (SAM system) to upregulate endogenous CD14 gene expression as an approach to stably engineer MSCs with novel functions ^74^. We observed BALB-MSCs overexpressing CD14 responded more quickly to LPS stimulation (as measured by NF-κB translocation) and displayed enhance antibacterial activity (as evident in inhibition of co-cultured *E. coli*). Surprisingly, we found that overexpression of TLR4 did not have the same effect. This suggests that Cd14 may serve as a bottleneck in controlling sensitivity to LPS stimulation in MSCs, similar to what has been observed other cell types ^73,79^. A recent study by Jiang et al. similarly concluded that CD14 deficiency in MSCs may be responsible for an attenuated LPS response ^82^. Our work builds on the framework outlined by Jiang et al. and demonstrates that the overexpression of CD14 in MSCs accelerates LPS response through increased NF-κB nuclear translocation rate and subsequent changes in global gene expression. LPS priming through TLR4 activation has been utilized to control diverse therapeutic properties in MSCs, including (but not limited to) hematopoietic support ^83–85^, immunomodulation ^40,86–88^, and antibacterial activity ^21,44^. A recent report by Munir et al. demonstrated that LPS-primed MSCs have enhanced therapeutic properties, as administration of these cells dramatically accelerated wound healing by reshaping the wound site through increasing recruitment and activation of neutrophils and macrophages ^89^. How might combining CD14 overexpression in MSCs with TLR4-stimulation impact these diverse therapeutic properties? Subsequent studies in this field may benefit from the use of CD14 overexpression as a mechanism to reduce the amount of LPS required for stimulation, or to amplify therapeutic effects. The role of CD14 in MSC biology has remained elusive because this protein, by early definitions, was considered a negative marker for this cell type ^46^. However, studies have since identified MSC subpopulations in mice and humans that express the CD14 protein, further strengthening the conclusion that MSCs are a heterogeneous cell population even within a single organism ^82,90^.

The emergence of novel single-cell characterization technologies allows for a higher resolution molecular understanding of MSC subpopulations. These tools are beginning to be used to define MSC heterogeneity, and to explore how genetic modifications and priming can impact MSC population diversity ^51,52,91,92^. Strategies to limit heterogeneity and enrich homogenous populations of MSCs are critical for ensuring safe and effective MSC-based therapies. We used 10X Genomics single-cell transcriptomics technology to determine the effect that CD14 expression and LPS priming have on MSC population heterogeneity. Interestingly, based on two methods of dimensionality reduction (tSNE and UMAP), C57-MSCs appear more homogenous than BALB-MSCs, which may partially explain why the latter have a more attenuated LPS response. Unexpectedly, we observed that overexpression of CD14 in BALB-MSCs not only increased the speed with which the cells responded to LPS, but also changed the transcriptional ground state of BALB-MSCs in the absence of LPS exposure. One possible explanation is that there are trace levels of LPS present in our culture system, not enough to stimulate wildtype BALB-MSCs but sufficient to activate signal transduction programs when recognition is facilitated by overexpression of CD14. Indeed, there is evidence that CD14 can increase sensitivity to LPS in a number of physiological contexts ^72,73,79,93^. Alternatively, CD14 may be recognizing a different ligand, such as Gram-positive cell wall components HSP60 or LAM, or other endogenous lipids present in culture ^81,94–97^. Regardless of the stimulating ligand, the expression of CD14 on the surface of MSCs makes them more responsive to LPS treatment, likely by transitioning these cells to a primed state.

Through our transcriptional profiling analyses, we observed that C57-MSCs and BALB-MSCs exhibited different expression of multiple TLRs. A notable example is that BALB-MSCs had higher baseline expression of TLR1, TLR2, and TLR6, which can form heterodimers (TLR1/2 and TLR2/6) to sense and respond to lipoprotein. In our LPS priming experiments, cells were treated with a standard LPS preparation (not an ultra-pure form) that includes remnants of other bacterial components including lipoproteins, allowing it to co-stimulate TLR2 in addition to TLR4. An elegant study by Kellogg et al. recently demonstrated that cells exhibit distinct NF-κB dynamics when stimulated by TLR2 vs TLR4 agonists ^98^. Additionally, when cells are exposed to a mixture of TLR2 and TLR4 stimulating agents, single cells do not exhibit a hybrid response and instead respond to one ligand or the other ^98^. Kellogg et al. demonstrated that cells signaling through TLR4 exhibit a rapid and unified response to LPS stimulation, while cells exposed to TLR2 ligands exhibit a delayed and variable response between individual cells ^98^. Using NF-κB nuclear translocation assays in our study, we observed the majority of C57-MSCs responded quickly to LPS treatment, which is consistent with the TLR4 signaling dynamics reported by Kellogg et al. In contrast, when wildtype BALB-MSCs were treated with LPS they exhibited delayed and longer duration NF-κB translocation kinetics, which is more reminiscent of the TLR2 dynamics. It is therefore possible that in response to LPS treatment or bacterial exposure, C57-MSCs respond through a TLR4 signaling program whereas BALB-MSCs respond through TLR2 signaling. Overexpression of CD14 in BALB-MSCs dramatically shifted their NF-κB nuclear translocation dynamics to a rapid response, similar to that of wildtype C57-MSCs. One interpretation of this result is that high levels of CD14 in MSCs increases their sensitivity to LPS treatment and promotes signaling through TLR4 rather than TLR2. This model is supported by the study from Sung et al. who demonstrated that TLR2 was not involved with antibacterial activity in MSCs, suggesting that TLR2-based signaling may be insufficient for direct inhibition of bacterial growth but may play a role in immunomodulation in response to infection ^21^. However, further experiments are needed to test this model by utilizing single-cell transcriptional profiles of CD14 expressing MSCs when exposed to TLR2 or TLR4 agonists.

A number of mechanisms may be at play when it comes to how MSCs limit bacterial growth. In previous studies, MSCs have been shown to exert antimicrobial pressure through secretion of bactericidal peptides LL-37 (Camp) and β-defensin 2 (Defb2) ^20,21^. We did not observe the MSCs used in this study to express Camp or Defb2 prior to or after LPS treatment, highlighting the extensive source-specific phenotypic variation of MSCs. There were, however, several notable genes with differential expression between MSC types and after LPS treatment that have established roles in limiting bacterial growth. In particular, Lipocalin-2 (Lcn2) is a bacteriostatic peptide that inhibits bacterial iron-sequestering siderophores and has previously been demonstrated to protect mice from *E. coli* induced pneumonia ^23,99^. We observed an increase in Lcn2 expression after LPS treatment in both C57-MSCs and BALB-MSCs, and C57-MSCs showed robust expression of Lcn2 prior to LPS treatment as well. Another notable candidate is nitric oxide synthase 2 (Nos2), which was also expressed during routine growth in C57-MSCs and was observed to increase in expression in both MSC types following LPS treatment. Nitric oxide secretion is one mechanism by which immune cells can limit microbial growth, and MSCs can stimulate macrophages to increase NO production which enhances their bacterial-killing properties ^100,101^. To our knowledge, Nos2 expression in MSCs has not previously been implicated in their antibacterial activity. Although we did not test these proteins in antibacterial assays in this work, these are examples of potential targets identified in this study that could be modulated using similar CRISPRa tools in subsequent antibacterial studies.

The work outlined in this study demonstrates another example of source-dependent phenotypes in MSCs. These source-specific phenotypic variations are a challenge towards the development of MSC-based therapies. How can we ensure that MSCs collected from different sources will exhibit the necessary therapeutic behaviors? This becomes increasingly problematic if autologous MSC transplants are used, requiring phenotypic characterization of each patient’s MSCs prior to administration. Additionally, MSCs are inherently heterogenous, even within populations that were isolated from a single source. How can we homogenize these cells and harness subpopulations that are enriched for desired properties? In this work we demonstrate that priming and targeted genetic modifications using CRISPR tools can drive MSCs towards specific therapeutic states. MSCs have immense therapeutic potential, however their progression into the clinic has been stymied by our lack of understanding of these cells due to their inconsistencies and heterogeneity. As we continue to learn how these unique cells operate and develop methods that effectively direct their behavior, the closer we get towards safe and reliable MSC-based therapies.

## MATERIALS AND METHODS

### Cell culture and media

Bone marrow derived MSCs from C57BL/6 and BALB/c mice were purchased commercially and routinely cultured using Mouse Mesenchymal Stem Cell Media (Cyagen US Inc.). Primary MEFs from C57BL/6 mice (Blelloch Lab, UCSF), primary MDFs from C57BL/6 mice (ScienCell Research Laboratories, Carlsbad, CA), NIH/3T3 cells (ATCC, Manassas, VA) and HEK 293T cells (ATCC) were routinely grown in high glucose, pyruvate DMEM (4.5 g/L D-glucose, 110 mg/L sodium pyruvate; Gibco™ Thermo Fisher Scientific, Waltham, Massachusetts) supplemented with 10% heat inactivated FBS (Gibco™ Thermo Fisher Scientific) and 1% penicillin-streptomycin. All mammalian cell culture was conducted in two-dimensional monolayers on tissue culture treated plastic dishes or well-plates in a humidified incubator at 37°C with 5% CO_2_. MSCs were used in experiments at a passage number ≤ 10 and subcultured when cells reached ~80-90% confluence. All LPS-priming experiments were conducted in RPMI 1640 media (Gibco™ Thermo Fisher Scientific) + 5% FBS without antibiotics using a standard preparation of LPS from *E. coli* 055:B5 (InvivoGen, San Diego, California) at a final concentration of 100 ng/ml.

### Differentiation Assays

Differentiation assays were performed using adipogenic and osteogenic differentiation media as per instructions provided (Cyagen Biosciences, Inc.). In brief, for adipogenic assays, cells were plated at 2×10^4^ cells/cm^2^ in a 6-well tissue culture treated dish in MSC basal media and grown until cells reached 100% confluence. Cells were then induced to undergo adipogenesis by culturing cells for 3 days in MSC Adipogenic Differentiation Induction Media (Cyagen Biosciences, Inc.), followed by 1 day in MSC Adipogenic Differentiation Maintenance Media (Cyagen Biosciences, Inc.). The process of cycling adipogenic induction and maintenance medias was repeated for a total of three times. Cells were then fixed using 4% paraformaldehyde in PBS for 30 minutes at room temperature, washed twice with PBS and stored at 4°C. Fixed cells were then stained with an Oil Red O solution (diluted 3:2 in DI water and filtered through filter paper) for 30 minutes and washed 3 times with PBS. For osteogenic differentiation assays, 6-well cell culture dishes were first coated with 0.1% gelatin solution for 30 minutes at room temperature and then gelatin solution was removed. Cells were seeded at 2×10^4^ cells/cm^2^ in MSC basal media and grown to ~70% confluence. Cells were then put in MSC Osteogenic Differentiation Media (Cyagen Biosciences, Inc.), and this media was replaced every 3 days for a total incubation of 2 weeks. After two weeks incubation in Osteogenic Differentiation Media, cells were fixed with 4% paraformaldehyde in PBS as described above. Fixed cells were then stained with 1 ml Alizarin Red S working solution (Cyagen Biosciences, Inc.) for 5 minutes at room temperature, and then washed 3 times with PBS. Oil red O and Alizarin Red S stained cells were imaged using phase contrast microscopy with an EVOS XL Core Cell Imaging System (Thermo Fisher Scientific).

### Immunostaining MSC surface markers and flow cytometry

Cells were fixed with 4% paraformaldehyde in PBS for 15 minutes at room temperature and then washed twice with PBS. Fixed cells were incubated in Blocking Buffer (3% BSA in PBS, filtered with 0.22 μm filter) at room temperature overnight. For mouse MSC surface markers, cells were stained using the Mouse Mesenchymal Marker Antibody Panel (R&D Systems). Primary antibodies were added to cells at a concentration of 1 μg/ml diluted in Blocking Buffer and incubated at room temperature for 1 hour on a rocker. Cells were then washed twice with PBS and resuspended in goat anti-rat IgG Alexa Flour 647 secondary antibody (Abcam, Cambridge, United Kingdom) diluted 1:1000 in Blocking Buffer and incubated at room temperature for 30 minutes. Cells were then washed and resuspended in PBS, and fluorescence levels of 10,000 cells were measured using a BD Accuri C6 Plus flow cytometer. Flow cytometry data was analyzed and plotted using FlowJo v10 (BD, Franklin Lakes, NJ).

### Antibacterial Assays

Protocols for antibacterial assays used in this study were adapted from previously described methods ^20^. In brief, MSCs were trypsinized, quenched with media, and then washed twice with PBS. Cells were then resuspended in RPMI + 5% FBS without antibiotics and cell density was quantified using a Bio-Rad TC20 automated cell counter. Cells were added to tissue-culture treated multi-well dishes at a concentration of 1.25 × 10^5^ cells / cm^2^ in RPMI + 5% FBS without antibiotics (+/− LPS 100 ng/ml) and incubated overnight at 37°C + 5% CO_2_. *E. coli* strain K-12 MG1655 was grown overnight in LB liquid at 37°C while shaking at ~220 RPM. The following day, *E. coli* cells were enumerated by OD_600_ measurement (1 OD = 4 × 10^8^ CFU/ml). Bacterial cells were then diluted in RPMI + 5% FBS and added directly to mammalian cell culture wells to a final concentration of 1 × 10^3^ CFU/ml (final volume of *E. coli* suspension added was ~2% of total incubation volume). Mammalian cells and *E. coli* were co-incubated at 37°C + 5% CO_2_ for 6 hours without agitation. After 6 hours, *E. coli* viability was measured by plating dilutions of bacterial cell suspensions onto LB agar and counting CFUs. Statistical significance between groups was determined using t-test and plotted using Prism 8 software (GraphPad). For antibacterial assays, we used the pwr.t.test function in R to calculate that given the CFU values we were obtaining to have an average power of 80% across conditions we needed n=5 per condition (some conditions have more). In addition, we used the Benjamini-Hochberg procedure which adjusted for multiple comparisons to determine that all p-values below 0.044 represent significant results.

### NF-κB nuclear translocation assays

MSCs were plated into black-walled 96-well glass bottom tissue culture plates in RPMI media + 5% FBS at a cell density of ~60,000 cells / cm^2^. Adherent cells were left untreated, or treated with LPS from *E. coli* 055:B5 (100 ng/ml) and collected at specific timepoints post-exposure (0.083 hr, 0.25 hr, 0.5 hr, 1 hr, 2 hrs, and 18 hrs). Media was removed from wells and cells were fixed with 4% paraformaldehyde in PBS for 15 minutes at room temperature, and then washed twice with PBS. The cells were then incubated in Blocking Buffer B (3% BSA, 3% normal goat serum, 0.1% Triton X-100 in PBS, filtered through 0.22 μm filter) for 1 hour at room temperature. NF-κB p65 (L8F6) Mouse mAb antibodies (Cell Signalling Technology, Danvers, MA) diluted in blocking buffer were then added to the fixed cells and incubated for 18 hours at 4°C. The samples were then washed 3 times with 0.1% Triton X-100 in PBS before incubation with goat anti-mouse IgG Alexa Fluor 488 secondary antibody (Abcam) for 1 hour at room temperature. The cells were then washed once with PBS and incubated with DAPI diluted in PBS for 10 minutes, followed by two more PBS washes. Images of the samples were acquired using the CellInsight CX7 High-Content Screening platform and analyzed using the HCS Studio Cell Analysis software (Thermo Fisher Scientific). Background subtraction was first performed, followed by image segmentation using the DAPI channel to count the total number of cells imaged in each field of view and record the average intensity values of the p65 channel for each nucleus. A ring region of interest (ROI) surrounding the nucleus for each cell was also created to measure antibody fluorescence in the cytoplasm. NF-κB p65 nuclear translocation was identified by counting cells yielding a ratio of nuclear average intensity to cytoplasmic intensity greater than 1. For nuclear translocation assays, we used the pwr.t.test function in R to calculate that given percent translocation values we were obtaining across time points, to have an average power of 80% across comparisons we needed n=3 per condition. In addition, we used the Benjamini-Hochberg procedure which adjusted for multiple comparisons to determine that all p-values below 0.00013716 represent significant results.

### Cytokine profiling

For cytokine analyses, ~250,000 cells were plated in 24-well tissue culture treated dishes and grown in 0.5 ml RPMI + 5% FBS and incubated at 37°C + 5% CO2 overnight. Cells were either left untreated, or treated with LPS from *E. coli* 055:B5 at a concentration of 100 ng/ml for ~18 hrs. Conditioned media was collected and centrifuged briefly to remove debris. Bead-based immunoassays were performed using LEGENDplex assays with the pre-defined Mouse Inflammation Panel and the Mouse Proinflammatory Chemokine Panel. LEGENDplex immunoassays were performed as described by the manufacturer’s protocols (BioLegend, San Diego, CA). LegendPlex assays were measured using a BD Accuri C6 Plus flow cytometer, and data analyzed using LEGENDplex software (BioLegend) and Prism 8 (GraphPad Software, San Diego, CA).

### Population level RNA-seq and data analysis

To harvest RNA from cells for RNA-seq, cells were resuspended in TRIzol reagent (Invitrogen™, Thermo Fisher Scientific) and stored at −80°C until further processing (n = 3 biological replicates per condition). For RNA-seq of cells during standard growth, RNA was collected from cells when they reached ~80-90% confluence. For the LPS-timecourse RNA-seq experiment, cells were plated at a concentration of 125,000 cells/cm^2^, treated with LPS (100 ng/ml) or without (samples labeled as 0 hr timepoint), and collected in TRIzol at 0.5 hr, 2 hr, and 18 hr post-exposure. RNA was purified from TRIzol using PureLink RNA Mini Kit (Invitrogen™, Thermo Fisher Scientific) and RNA concentration was measured with a Qubit fluorometer (Thermo Fisher Scientific). The quality of RNA preps was determined using the RNA 6000 Nano Kit on a Bioanalyzer (Agilent Technologies, Santa Clara, CA) and all samples used in this study had RNA integrity numbers > 7. RNA-seq libraries were generated using the KAPA HyperPrep Kit with RiboErase (HMR) and KAPA dual-indexed adapters (Roche Sequencing and Life Sciences, Wilmington, MA). RNA-seq libraries were quantified using the High Sensitivity DNA Chip on a Bioanalyzer (Agilent Technologies, Santa Clara, CA) and KAPA library quantification kit (Roche). RNA-seq libraries were sequenced using the Illumina NextSeq 500/550 platform with the High Output v2 kit (150 cycles) on paired-end mode (Illumina Inc, San Diego, CA). BCL files were converted to FASTQ and demultiplexed using the bcl2fastq conversion software (Illumina, Inc.). Quality filtering and adaptor trimming were performed using fastp ^102^ with following parameters: --qualified_quality_phred 25, -- cut_window_size 5 −3 and --cut_mean_quality 25. Kallisto was used for alignment-free mapping with 100 bootstraps per sample to calculate transcript counts ^103^. PCA was performed using the prcomp function in R, and MDS performed with edgeR ^104^. Differential expression analysis was performed using DeSeq2 ^105^ with significant genes being defined with a fold change > 2 and an adjusted p-value < 0.05. Pathway analysis was performed and visualized using Pathview ^71^. GO-term analysis was performed using PANTHER ^106,107^ and visualizations were plotted in R.

### Quantitative membrane proteomics

Cells were prepared using standard growth conditions as described above and processed when cell density reached ~75% - 95% confluency. Adherent cells were washed twice with DPBS and detached from the surface of the plate using a cell scraper into centrifuge tubes. Cells were centrifuged for 5 minutes at 300 RCF, supernatant removed, and cell pellets were flash frozen in liquid nitrogen. Each cell type was collected in biological triplicate and sent for membrane processing, protein extraction and LC-MS/MS (performed by MS Bioworks, Ann Arbor, Michigan). Cell pellets were processed using the Pierce MEM-PER reagent according to the manufacturers protocol. Extracted proteins were concentrated by trichloroacetic acid, and protein pellets were washed with ice cold acetone and solubilized in 8M Urea, 150mM NaCl, 50mM Tris-HCl pH8, 1X Roche Complete protease inhibitor. The protein concentration of the extract was determined by Qubit fluorometry. 25μg of protein was reduced with dithiothreitol, alkylated with iodoacetamide and digested overnight with trypsin (Promega). The digestion was terminated with formic acid and desalted using an Empore SD solid phase extraction plate. 2μg of each peptide sample was analyzed by nano LC-MS/MS with a Waters NanoAcquity HPLC system interfaced to a ThermoFisher Q Exactive. Peptides were loaded on a trapping column and eluted over a 75μm analytical column at 350nL/min; both columns were packed with Luna C18 resin (Phenomenex). The mass spectrometer was operated in data-dependent mode, with the Orbitrap operating at 70,000 FWHM and 17,500 FWHM for MS and MS/MS respectively. The fifteen most abundant ions were selected for MS/MS. 4hrs of instrument time was used for the analysis of each digest. Data were processed with MaxQuant version 1.6.0.13 ^108^. Differential protein expression analysis and clustering was performed using the DEP R package ^109^ to identify and characterize protein composition of cell membranes. GO-term analysis of membrane proteins was performed using PANTHER ^106^.

### CRISPR-mediated gene activation in MSCs

Gene activation by CRISPR was performed using the CRISPR/Cas9 Synergistic Activation Mediator (SAM) system ^74^. C57-MSC-CRISPRa and BALB-MSC-CRISPRa were constructed through lentiviral transduction of vectors lenti-dCas9-VP64_Blast (Addgene # 61425) and lenti-MS2-P65-HSF1_Hygro (Addgene #61426). Mouse CD14 and TLR4 sgRNAs sequences were designed using the sgRNA design tool from the Zhang Lab (http://sam.genome-engineering.org/database/). Six sgRNAs were designed per gene with different distances upstream of the transcription start site. Primers for individual sgRNAs were annealed and cloned into the lenti-sgRNA(MS2)_zeo backbone (Addgene #61427) using the Golden Gate cloning reaction into the BsmBI site (For primers sequences see **Supplementary Table 1**). sgRNA plasmids were checked for correct sequence integration using Sanger Sequencing (Genewiz Inc, South Plainfield, NJ). Each sgRNA lentiviral construct was transfected into HEK 293T cells along with packaging vectors psPAX2 (Addgene #12260) and pCMV-VSV-G (Addgene #8454) using Lipofectamine 3000 (Thermo Fisher Scientific), and incubated for 48 hours. HEK 293T media containing lentivirus was then filtered through a 0.45 μm syringe filter, added to MSCs with polybrene (10 μg/ml), and incubated for 24 hours. Following 24-hour transduction, the media was replaced with fresh media with appropriate antibiotic selection. Following selection, expression levels of CD14 and TLR4 were tested in CRISPRa cells expressing different sgRNAs using immunostaining followed by flow cytometry. Cells were live-stained using PE anti-mouse CD14 antibody or APC anti-mouse TLR4 antibody (BioLegend), then fixed with 4% paraformaldehyde for 15 minutes, washed and diluted in PBS. Stained cells were then examined using a BD FACSMelody™ Cell Sorter (BD Biosciences) and data was analyzed using FlowJo v10.

### Single-cell RNA-seq sample preparation

The cells used in scRNA-seq studies were C57-MSC-CRISPRa-Scrambled (expressing scrambled sgRNA-1), BALB-MSC-CRISPRa-Scrambled (expressing scrambled sgRNA-1), and BALB-MSC-CRISPRa-CD14 (expressing CD14-83 sgRNA). All cells were plated at a density of 125,000 cells/cm in 24-well plates in RPMI + 5% FBS and either treated with LPS (100 ng/ml for 0.5 hr, 2 hr, and 18 hr) or left untreated. Cells were then trypsinized, washed with PBS, and resuspended in RPMI + 5% FBS at a final concentration of 1×10^6^ cells/ml. Cell suspensions were then processed to isolate ~2,000 individual cells per sample using the Chromium Single Cell 3’ Reagent Kit v3 and 10x Chromium Controller following protocols as outlined in the corresponding 10x user guide Rev. B (10x Genomics, Pleasanton, CA). All subsequent steps (reverse transcription, cDNA amplification, and gene expression library construction) were performed as described in the 10x Chromium Single Cell 3’ Reagent Kits v3 Rev B User Guide (10x Genomics). Final libraries were quantified using a Qubit Fluorometer with hsDNA reagent (Thermo Fisher Scientific) and run on a Bioanalyzer High Sensitivity DNA chip (Agilent Technologies). Libraries were sequenced using an Illumina NextSeq 500/550 platform with the High Output v2.5 kit with 26×98 paired-end reads. Sequencing output was processed using Cell Ranger software protocols (10x Genomics). In brief, BCL files were demultiplexed and converted to fastq using Cell Ranger mkfastq. Fastq files were processed for alignment, filtering, barcode counting, and UMI counting using the Cell Ranger “count” feature, and pooled by each GEM using the Cell Ranger aggr pipeline.

### Single-cell RNA-seq analysis

Seurat and Monocle3 R packages were used to process, filter and normalize single-cell RNA-seq data ^110–115^, and figures for tSNE, U-MAP and violin plots were generated by modifying standard pipelines. Briefly, samples were adjusted for cell cycle stage and filtered as follows: number of Features above 200, and percentage RPS and RPL < 12. Samples were then log-normalized and scaled. Data processed across two different experimental runs was normalized using COMBAT ^116,117^. The UMAP settings were reduction = “pca”, dims = 1:10, n.neighbors = 10, n.components = 10, min.dist = 0.05, local connectivity = 5. The tSNE settings were dims = 1:10, dims.use = 1:10, reduction.use=“pca”. All code provided upon request. Euclidian distance between points was calculated using 10 dimensions (for both tSNE and UMAP). Differentially expressed genes were identified using the FindMarkers module in Seurat. Gene ontology analysis on differentially expressed genes was performed using PANTHER ^106^.

## ACKNOWLEDGEMENTS

We would like to thank Joe Schoeniger and Ramdane Harouaka for review of the manuscript. We thank Colleen Courtney for advice and training using the CellInsight CX7. We thank Paul Scott and Adam Bemis from 10x Genomics for their help with the single cell sequencing using the 10x Chromium. We thank the Blelloch Lab at UCSF for providing MEFs used in this study. This work was supported by the Laboratory Directed Research and Development program at Sandia National Laboratories, a multi-mission laboratory managed and operated by National Technology and Engineering Solutions of Sandia, LLC, a wholly owned subsidiary of Honeywell International, Inc., for the U.S. Department of Energy’s National Nuclear Security Administration under contract DE-NA0003525.

## COMPETING INTERESTS

The authors do not have any competing interests to report.

## DATA AVAILABILITY

RNA sequencing data was deposited into the National Center for Biotechnology Information Sequence Read Archive (https://www.ncbi.nlm.nih.gov/sra) and is available under the study accession PRJNA667557.

## Notes

### Competing Interest Statement

The authors have declared no competing interest.

