## Supplemental material for "Augmentation of Antibacterial Activity in Mesenchymal Stromal Cells Through Systems-Level Analysis and CRISPR-mediated Activation of CD14"

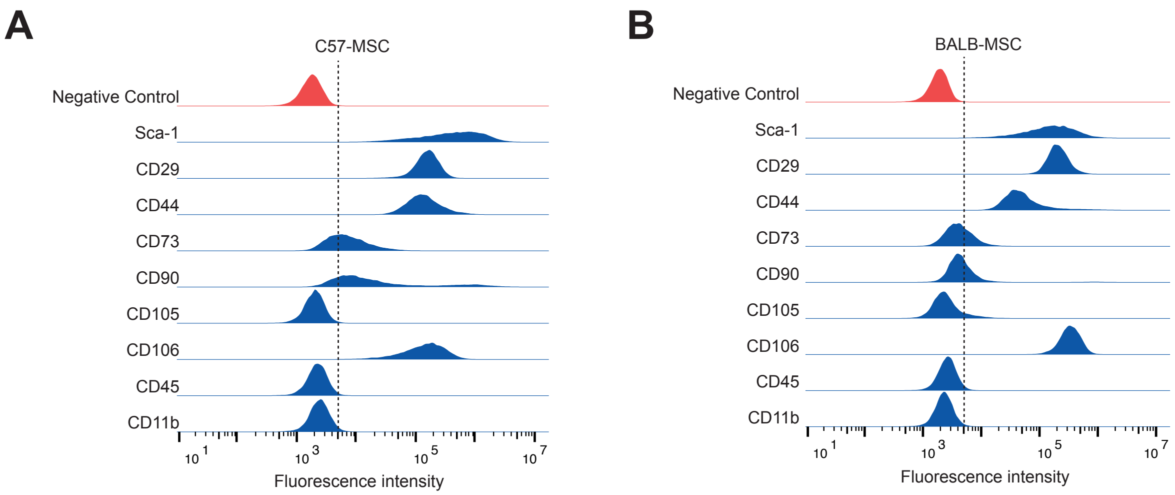


**Supplementary Figure 1. Analysis of traditional MSC cell-surface markers in different MSC types.** Cells were inspected for the expression of MSC cell-surface markers by immunostaining with the Mouse Mesenchymal Marker Antibody Panel (R&D Systems) followed by flow cytometry. Both **(A)** C57-MSCs and **(B)** BALB-MSCs stained positively (>99% of cells) for the MSC markers Sca-1, CD29, CD44 and CD106. The MSC markers CD73 and CD90 were observed only in subsets of cells analyzed indicating population heterogeneity that has been observed with these proteins in MSCs [1,2]. C57-MSCs and BALB-MSCs exhibited negative or low staining of CD105, respectively, and subpopulations of CD105-negative MSCs have also been previously characterized [3]. The negative MSC markers, CD45 and CD11b, were not detected in either C57-MSC or BALB-MSCs.

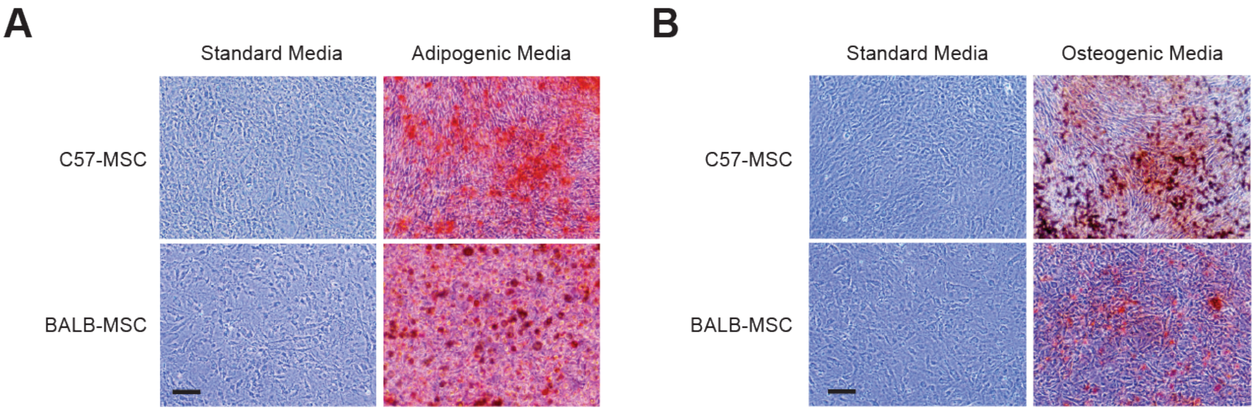


**Supplementary Figure 2. MSC differentiation assays.** C57-MSCs and BALB-MSCs were tested for their ability to differentiate using **(A)** adipogenic and **(B)** osteogenic assays (Cyagen, Inc). Using this approach, we observed both cell types to be capable of adipogenesis and osteogensis as visualized by Oil Red O and Alizarin Red S staining, respectively (Scale bars = 200 um).

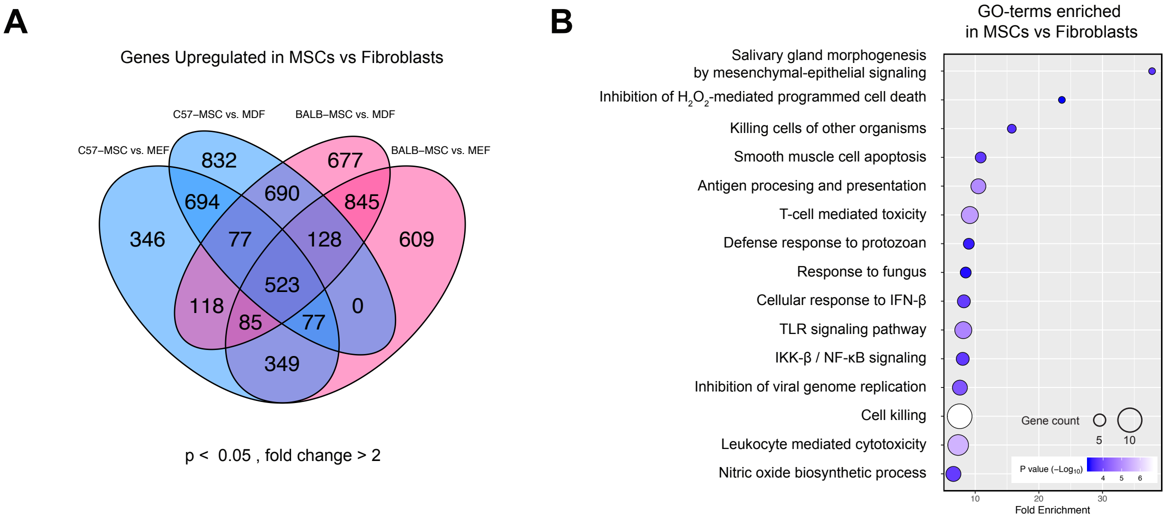


**Supplementary Figure 3. Transcriptional differences between MSCs and fibroblasts.** **(A)** Venn diagram showing numbers of genes upregulated in pairwise comparisons of MSCs vs fibroblasts (upregulated genes were defined as having a fold change > 2 and an adjusted P < 0.05). The center overlapping area of 523 genes represented shared upregulated genes between both MSC types when compared to both fibroblasts. **(B)** These MSC-specific genes were then inspected for functional enrichment using GO-term analysis. The most functionally enriched GO category involved the regulation of branching involved in salivary gland morphogenesis by mesenchymal-epithelial signaling (GO:0060665), which was a result of upregulation of Fgf7, Hgf, and Met. Of particular note, MSCs were enriched for the TLR-signaling pathway (GO:0002224) that included higher expression of genes involved with responding to bacterial PAMPs including Lbp, Tlr2 and Irf1. Additionally, MSCs were enriched for the genes involved with cell killing (GO:0031341) which included chemokines (Cxcl1 and Cxcl5), MHC genes (H2-BI, H2-T22, H2-Q6, H2-Q2, H2-K1), as well as pro-apoptotic genes (Stat5a, Bcl2l11, Gapdh, Arrb2).

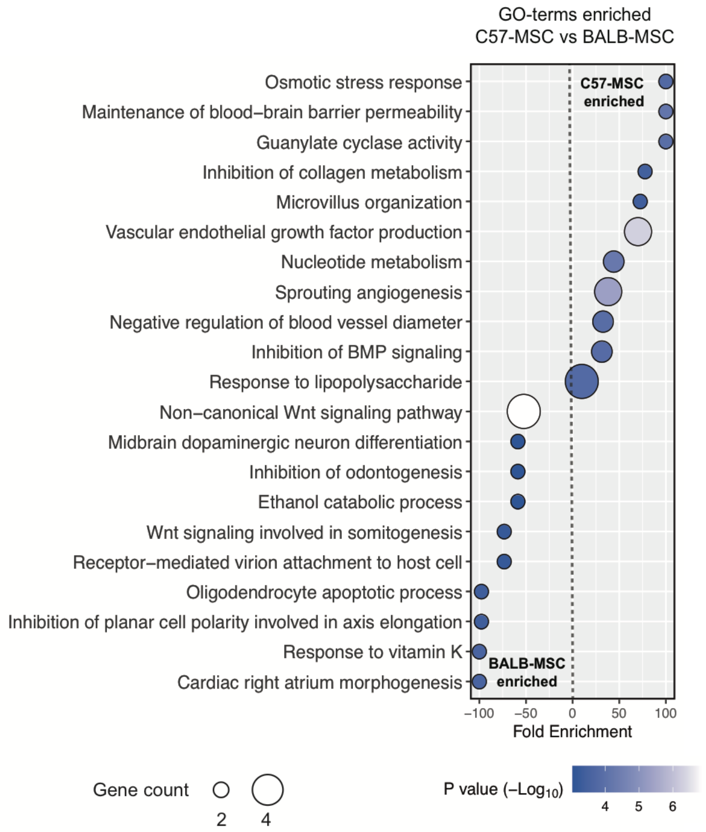


**Supplementary Figure 4. Comparative GO-term enrichment of C57-MSCs and BALB-MSCs.** RNA-seq data from C57-MSCs and BALB-MSCs were examined for differential gene expression using DeSeq2 [4]. Genes that were upregulated in C57-MSCs compared to BALB-MSCs (fold change > 2, adjust P < 0.05) were inputted into the PANTHER GO Enrichment Analysis to find biological processes that were enriched in C57-MSCs. The same process was also performed to identify GO-terms associated with genes upregulated in BALB-MSCs.

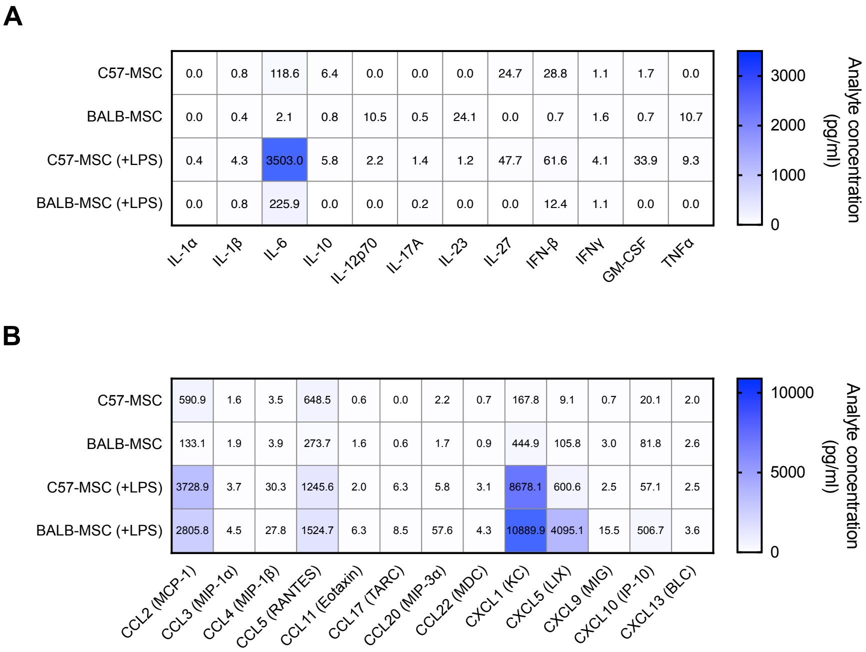


**Supplementary Figure 5. Cytokine profiling from MSCs with and without LPS-exposure.** Media was collected from C57-MSCs and BALB-MSCs with either no treatment, or after 18 hours of LPS-exposure and cytokine profiles were examined using LEGENDplex bead-based immunoassays. **(A)** Both cell types exhibited similar cytokine profiles with many of the cytokines assayed being low, or below the limit of detection. The most significant difference observed between cell types was the production of IL-6, with levels being ~60 fold higher in C57-MSCs (p=0.0079, t-test). IL-6 levels remained elevated in C57-MSCs after LPS-exposure and exhibited ~16-fold more IL-6 compared to BALB-MSCs. **(B)** Proinflammatory chemokines were also tested, and similar to the inflammation panel, most of the chemokines in the panel were not detected or were observed at low levels. CCL2 (MCP-1) was ~4 fold higher in C57-MSCs compared to BALB-MSCs (p=0.0042, t-test). After LPS treatment, both MSC types exhibited high levels of CCL2, CCL5 and CXCL1 (> 1 ng/ml), and we observed ~7-fold higher levels of CXCL5 produced by BALB-MSCs compared to C57-MSCs.

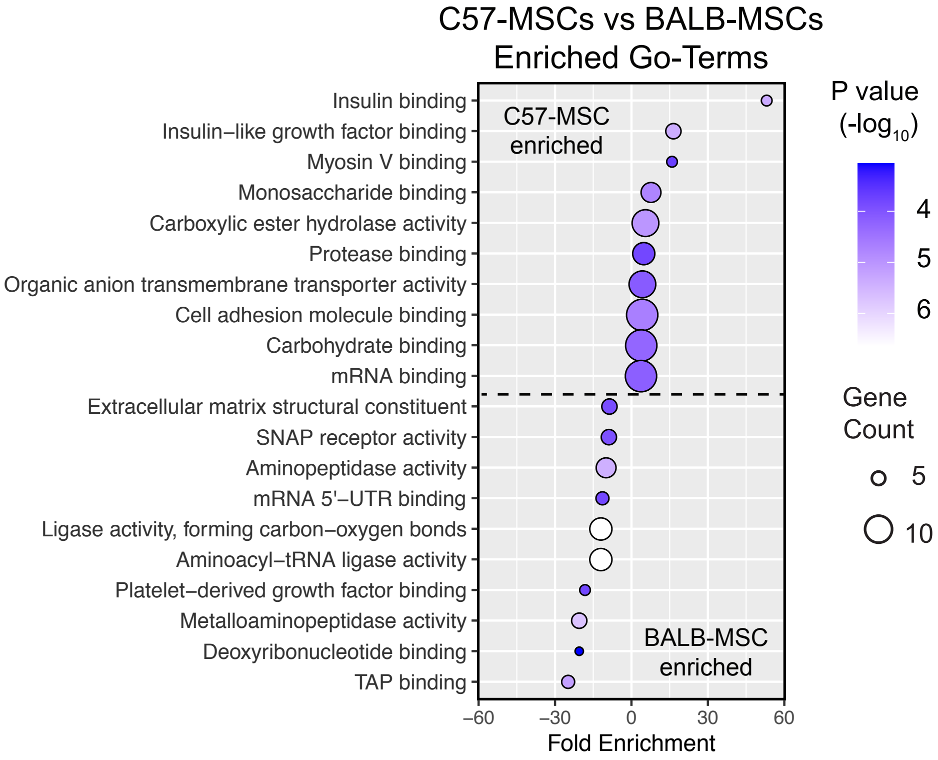


**Supplementary Figure 6. GO-term enrichment of membrane proteins between MSCs.** MSC membrane proteins were quantified using LC-MS/MS and analyzed for differential protein expression using the DEP R package [5]. Proteins upregulated in C57-MSC/BALB-MSC or BALB-MSC/C57-MSC were examined for enriched biological functions using GO-term analysis.

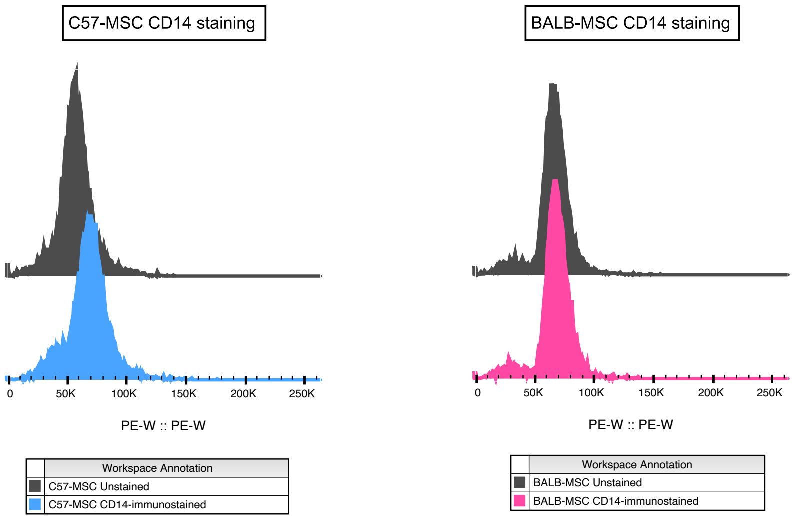


**Supplementary Figure 7. CD14 immunostaining in C57-MSCs and BALB-MSCs.** Both MSC types were live-stained using a PE conjugated anti-mouse CD14 antibody and expression levels were measured using flow cytometry. The mean fluorescence intensity (MFI) of immunostained MSCs was compared to MFIs from respective unstained controls. Using this approach, we observed increased fluorescence intensity of C57-MSCs stained with CD14 when compared to their unstained controls (MFI = 68198 vs. 57669, respectively), indicating the presence of CD14 protein expression. Alternatively, the BALB-MSCs stained with CD14 were more similar to their unstained controls (MFI = 67100 vs. 65679, respectively), indicating these cells have extremely low or absent CD14 expression.

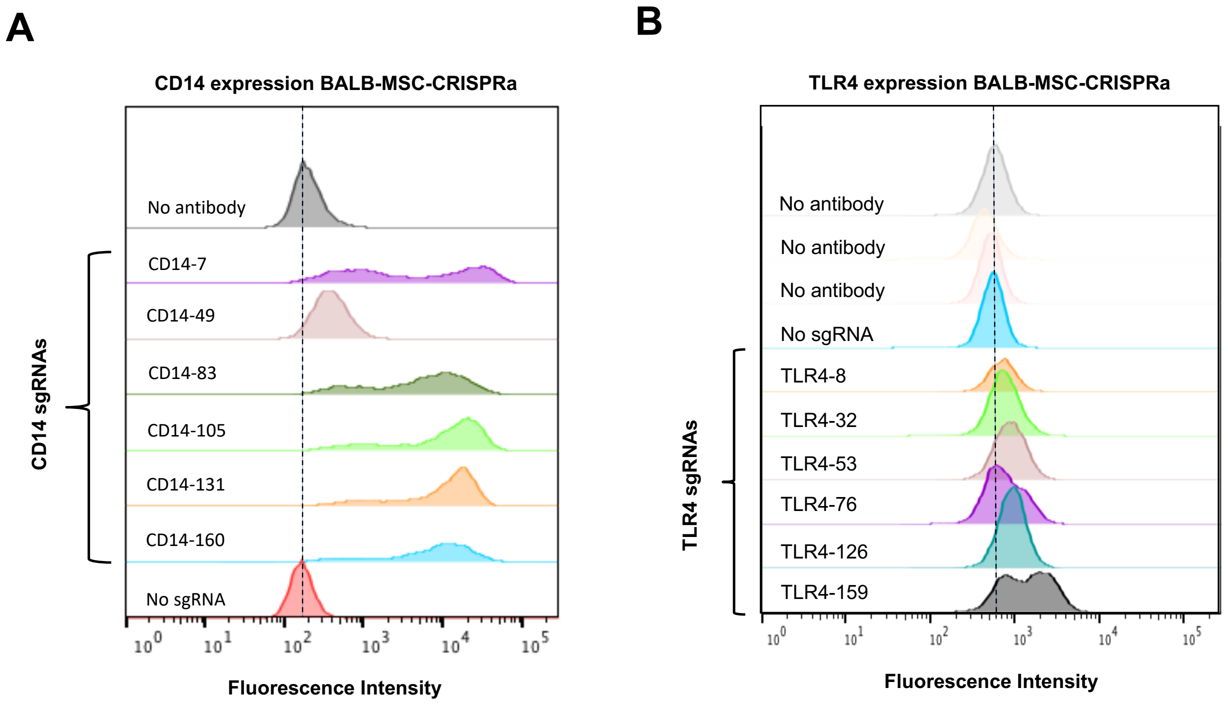


**Supplementary Figure 8. Comparison of sgRNAs to activate protein expression in BALB-MSC-CRISPRa cells.** BALB-MSCs expressing the CRISPRa SAM system were transduced with lentiviral constructs to express one of six sgRNAs that target either **(A)** CD14 of **(B)** TLR4. Each sgRNA was designed to target a different distance upstream of the transcription start site (represented by the number after the gene name). MSCs were live-stained using anti-CD14-PE or anti-TLR4-APC antibodies and examined for fluorescence intensity using flow cytometry. Negative controls used here were BALB-MSC-CRISPRa not expressing sgRNA with no antibody staining (labeled no antibody), or the same cells treated with antibody (labeled no sgRNA).

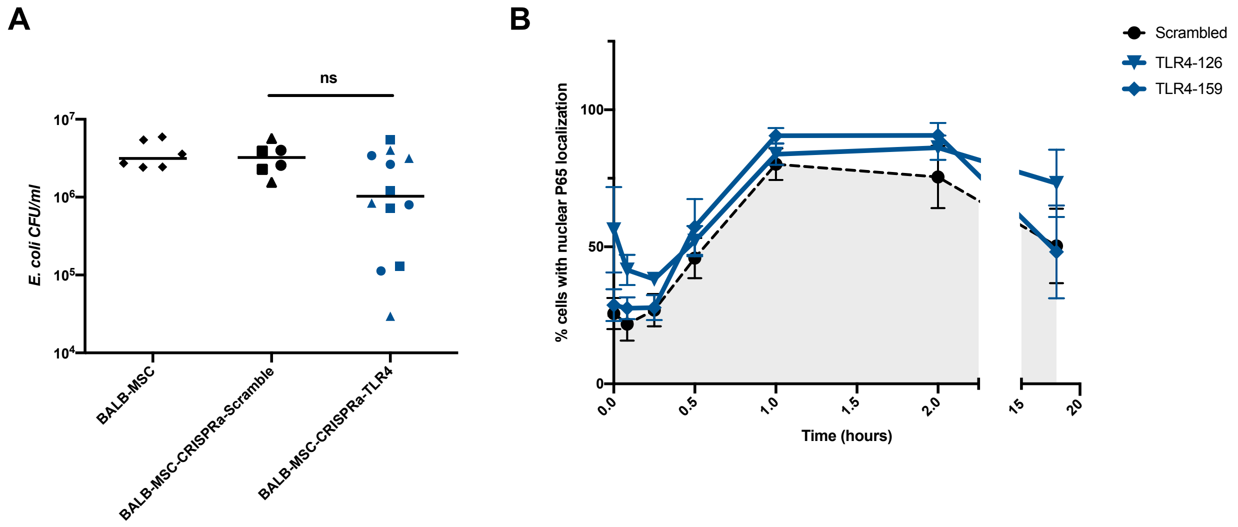


**Supplementary Figure 9. Functional analysis of TLR4 overexpression in BALB-MSCs.** **(A)** BALB-MSCs overexpressing TLR4 were examined for their antibacterial properties by co-culturing these MSCs with *E. coli* for 6 hours and measuring the CFUs in the media (P = 0.11, t-test). For scrambled sgRNAs: circle = non-targeting_control_1, square = non-targeting_control_2, triangle = non-targeting_control_3; For TLR4 sgRNAs: circle = TLR4-53, square = TLR4-126, triangle = TLR4-159. **(B)** BALB-MSC-CRISPRa-TLR4 cells were also inspected for their rates of NF-κB nuclear translocation after LPS-exposure compared to cells expressing a non-targeting “Scrambled” control.

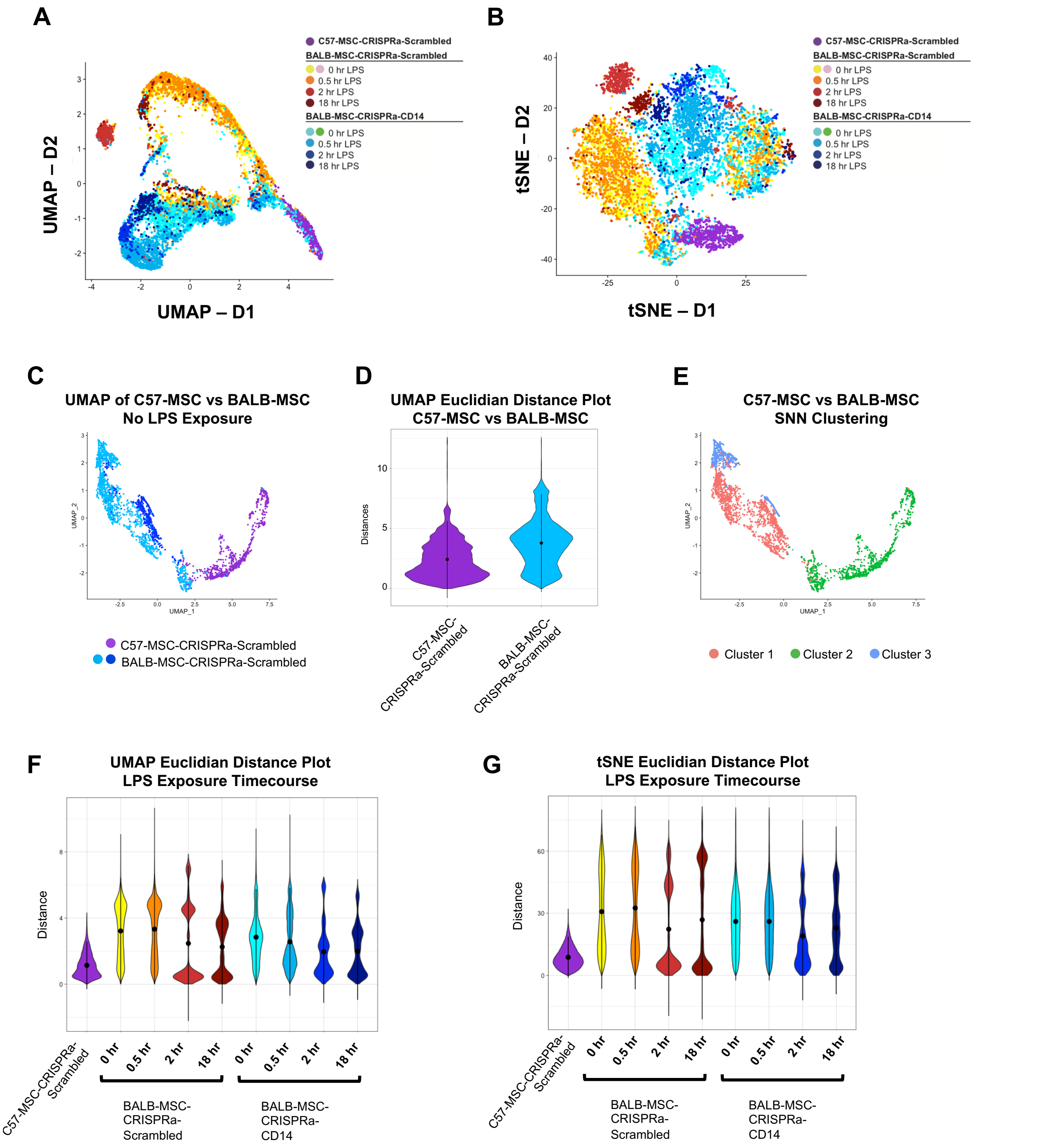


**Supplementary Figure 10. Single-cell RNA-seq visualization and Euclidean distance plots.** Population structure visualization of single-cell RNA-seq profiles from MSCs during LPS-exposure time course using **(A)** UMAP and **(B)** tSNE dimensionality reduction methods. **(C)** UMAP plot depicting single cell transcriptional profiles of untreated C57-MSC-CRISPRa-Scrambled and BALB-MSC-CRISPRa-Scrambled cells to examine transcriptional heterogeneity between cell types. **(D)** Pairwise Euclidian distance plot depicting distance measurements between all cells in the UMAP projection from panel C within each cell type. A Kruskal-Wallis non-parametric rank sum test shows a significance of $p < 2.2e-16$ with a chi-squared statistic of 141984, suggesting that C57-MSCs have decreased transcriptional heterogeneity compared to BALB-MSCs. **(E)** Seurat's shared nearest neighbors (SNN) algorithm was used to identify clusters of cells. Using a resolution value set at 0.1 we observed all C57-MSCs clustered together and BALB-MSCs were spread across three clusters (with some cells overlapping with the C57-MSC cluster). Cellular transcriptional heterogeneity during LPS-exposure were also measured using the pairwise n-dimensional Euclidian distances between members of the same experimental sample for both **(F)** UMAP **(G)** tSNE approaches.

| Primer Name | Sequence |
|---|---|
| CD14-7-F | CACCGGTACGCACCAGACAAGTCCG |
| CD14-7-R | AAACCGGACTTGTCTGGTGCGTACC |
| CD14-49-F | CACCGGAATAATGATCTAAGGCACT |
| CD14-49-R | AAACAGTGCCTTAGATCATTATTCC |
| CD14-83-F | CACCGGAAAATGGAGGTGAATCAAT |
| CD14-83-R | AAACATTGATTCACCTCCATTTTCC |
| CD14-105-F | CACCGTTGCTAGCAACTAAGACTAG |
| CD14-105-R | AAACCTAGTCTTAGTTGCTAGCAAC |
| CD14-131-F | CACCGAAGAGCTGGATTTGAACGGT |
| CD14-131-R | AAACACCGTTCAAATCCAGCTCTTC |
| CD14-160-F | CACCGTGAATGTAATTGGACATTTG |
| CD14-160-R | AAACCAAATGTCCAATTACATTCAC |
| TLR4-8-F | CACCGCAGATCGTCATGTTCTCTCA |
| TLR4-8-R | AAACTGAGAGAACATGACGATCTGC |
| TLR4-32-F | CACCGTGGTGGCAGCGCAGAGTCCC |
| TLR4-32-R | AAACGGGACTCTGCGCTGCCACCAC |
| TLR4-53-F | CACCGAGGGAAGAGGCAGGTGTCCC |
| TLR4-53-R | AAACGGGACACCTGCCTCTTCCCTC |
| TLR4-76-F | CACCGCTTGCAGAGGGGCACCCACT |
| TLR4-76-R | AAACAGTGGGTGCCCCTCTGCAAGC |
| TLR4-126-F | CACCGAACCTTAGCATTCTCACTTT |
| TLR4-126-R | AAACAAAGTGAGAATGCTAAGGTTC |
| TLR4-159-F | CACCGGAATCGATCTGCCCCGTCGC |
| TLR4-159-R | AAACGCGACGGGGCAGATCGATTCC |
| NonTargetingControl_0001_F | CACCGGCGAGGTATTCGGCTCCGCG |
| NonTargetingControl_0001_R | AAACCGCGGAGCCGAATACCTCGCC |
| NonTargetingControl_0002_F | CACCGGCTTTCACGGAGGTTCGACG |
| NonTargetingControl_0002_R | AAACCGTCGAACCTCCGTGAAAGCC |

| | |
|---|---|
| NonTargetingControl_0003_F | CACCGATGTTGCAGTTCGGCTCGAT |

**Supplementary Table 1. Primers used in this study.**
